# Peptide ligands for the affinity purification of adeno-associated viruses from HEK 293 cell lysates

**DOI:** 10.1101/2023.02.19.529155

**Authors:** Wenning Chu, Shriarjun Shastry, Eduardo Barbieri, Raphael Prodromou, Paul Greback-Clarke, Will Smith, Brandyn Moore, Ryan Kilgore, Christopher Cummings, Jennifer Pancorbo, Gary Gilleskie, Michael A. Daniele, Stefano Menegatti

**Affiliations:** Department of Chemical and Biomolecular Engineering, North Carolina State University, 911 Partners Way, Raleigh, NC 27606, USA; Biomanufacturing Training and Education Center (BTEC), North Carolina State University, 850 Oval Dr, Raleigh, NC 27606, USA; North Carolina Viral Vector Initiative in Research and Learning (NC-VVIRAL), North Carolina State University, 911 Oval Dr, Raleigh, NC 27695, USA; Joint Department of Biomedical Engineering, North Carolina State University and University of North Carolina at Chapel Hill, 911 Oval Drive, Raleigh, NC 27695, USA; LigaTrap Technologies LLC, Raleigh, NC 27606

**Keywords:** Adeno-associated virus, peptide ligands, affinity chromatography, virus purification, HEK 293 lysate

## Abstract

Adeno-associated viruses (AAVs) are the vector of choice for delivering gene therapies that can cure inherited and acquired diseases. Clinical research on various AAV serotypes significantly increased in recent years alongside regulatory approvals of AAV-based therapies. The current AAV purification platform hinges on the capture step, for which several affinity resins are commercially available. These adsorbents rely on protein ligands – typically camelid antibodies – that provide high binding capacity and selectivity, but suffer from low biochemical stability and high cost, and impose harsh elution conditions (pH < 3) that can harm the transduction activity of recovered AAVs. Addressing these challenges, this study introduces peptide ligands that selectively capture AAVs and release them under mild conditions (pH 6.0). The peptide sequences were identified by screening a focused library and modeled *in silico* against AAV serotypes 2 and 9 (AAV2 and AAV9) to select candidate ligands that target homologous sites at the interface of the VP1-VP2 and VP2-VP3 virion proteins with mild binding strength (K_D_ ∼ 10^-5^-10^-6^ M). Selected peptides were conjugated to Toyopearl resin and evaluated via binding studies against AAV2 and AAV9, demonstrating the ability to target both serotypes with values of dynamic binding capacity (DBC_10%_ > 10^13^ vp per mL of resin) and product yields (∼50-80%) on par with commercial adsorbents. The peptide-based adsorbents were finally utilized to purify AAV2 from a HEK 293 cell lysate, affording high recovery (50-80%), 80-to-400-fold reduction of host cell proteins (HCPs), and high transduction activity (up to 80%) of the purified viruses.

## 1. Introduction

Gene therapy provides a unique approach to cure inherited and acquired diseases by downregulating or replacing a defective gene with a functional one^1,2^. As of 2022, almost 3,000 gene therapy clinical trials have been initiated^3,4^ and four gene therapy products approved by the Food and Drug Administration (FDA)^3–5^. A key role in the gene therapy revolution is played by viral vectors, owing to their ability to deliver a genomic payload efficiently and selectively to a target cell or tissue. While several classes of viral vectors are known and utilized today – employed in cell engineering (*e.g.*, Lentivirus and Baculovirus)^6–8^ or vaccination and oncolytic applications (*e.g.*, Adenovirus and Herpes Simplex Virus) ^9–11^ – the field of gene therapy is dominated by Adeno-Associated Viruses (AAV) owing to their low toxicity/pathogenicity and efficient integration of the transgene into the host cells.

AAV is a small, non-enveloped icosahedral virus, whose capsid can pack a linear single-strand DNA (ssDNA) genome of up to about 5 kilobases^18,19,20,21^. AAV capsids are formed by three virion proteins (VP1, VP2, and VP3), typically assembled in a 1:1:10 ratio^22^. To date, 13 distinct AAV serotypes (AAV1 - AAV13) are known – with the AAV2 being the most studied^23^ – which share a 65-99% sequence identity in their VPs and a 95-99% structural identity^1,24^. The biomolecular variations among serotypes translate in specific cell/tissue tropism: cardiac, skeletal, and muscle cells are targeted by AAV1, AAV6, and AAV9; retina cells by AAV2 and AAV8; hepatocytes by AAV8, AAV9, and AAV-DJ; lung cells by AAV5, AAV6, and AAV9; cells in the central nervous system (CNS) by AAV1, AAV5, AAV6, AAV9, and AAV-rh10^13,25–27^. Recently, recombinant AAVs (rAAVs) have been introduced, which feature improved gene packing and tissue tropism as well as lower immunogenicity and hepatotoxicity^28^.

The manufacturing of AAVs relies on two expression systems, namely triple transfected human embryonic kidney (HEK 293) cells, which generate ∼10^14^ vector genome-containing particles (vg) per liter of cell culture when harvested just 72 hours post-transfection and are ideal for serving small cohort of patients, such as those suffering from rare diseases^29^; and the live baculovirus infection of *Spodoptera frugiperda* (Sf9) insect cells, which can be grown in serum-free media and avoid the replication of contaminating human agents, and are ideal for large AAV batches^30^, such as those dedicated to fighting cancer and specific monogenic diseases^3,31^.

The large number of particles needed for a single patient dosing, which can reach up to 10^14^ vg per kg of body weight^32,33^, combined with stringent requirements of purity puts significant pressure on the downstream segment of the manufacturing process. The current platform process for AAV purification – reminiscent of the one established for mAbs – begins with an affinity-based capture step, which is tasked with removing most of the host cell proteins (HCPs) and DNA (hcDNA), and concentrating the AAV product for the subsequent steps of polishing and enrichment of full capsids^13,34,35^. Current affinity adsorbents include chromatographic resins functionalized with heparin^36,37^, whose applicability is limited to AAV2^38–40^, or camelid single-domain antibodies^41^. The latter include AVB Sepharose™ High Performance resin, which targets AAV serotypes 1, 2, 3, and 5^42^; the POROS™ CaptureSelect™ AAVX affinity resin, which targets AAV1 – AAV8, AAVrh10, and rAAVs^43,44^; POROS™ CaptureSelect AAV8 and AAV9 resins, specific to AAV8 and AAV9^43,45,46^; and AVIPure^®^ AAV2, AAV8, and AAV9 affinity resins^47^. Despite their excellent binding capacity (> 10^13^ vp per mL of resin) and selectivity^48,49^, these adsorbents feature high cost, low biochemical stability and short lifetime (< 20 cycles), and require harsh elution conditions (pH < 3.0) that can cause denaturation and aggregation of the AAV capsids, with consequent loss of transduction activity of the product^42–44,46^.

In seeking robust alternatives to protein ligands, we developed an ensemble of synthetic peptides that bind AAVs selectively, enable their elution under near-physiological conditions, and can be reused multiple times without losing binding strength and selectivity. The peptide ligands presented in this work target conserved binding sites found in all AAV serotypes *(i)* via multisite interactions, which provide the necessary binding strength and capacity for effective product capture, although *(ii)* the single AAV:peptide complexes can be easily dissociated, thus enabling high product recovery at mild elution conditions.

## 2. Results and Discussion

### 2.1. Selection of AAV-targeting ligands via rational design and screening of combinatorial peptide libraries

The biomolecular features that differentiate the various AAV serotypes – namely, the amino acid sequences of the virion proteins VP1, VP2, and VP3, and their unique arrangement within the capsid – also determine their behavior in terms of tissue tropism, transduction efficiency, and patient safety^74,75^. Key domains are displayed on the protrusions found on the fivefold cylinder and on the lines drawn between two contiguous threefold axes, and the twofold and fivefold axes^1^. As active sites, however, these domains are not suitable binding targets, since the association and dissociation with affinity ligands may cause structural and biochemical alterations leading to unwanted loss of tissue tropisms and transduction efficiency. Conversely, highly conserved regions (CRs) are found in all serotypes’ capsids, including a core eight-stranded β-barrel motif (βΒ-βI) and a α-helix (αA)^1,76,77^ on the convex side of the VPs (**Figure S1A**), that are not implicated in receptor binding, transduction, and antigenic specificity. These represent ideal target regions to universal AAV-binding peptides serving as ligands for serotype-independent purification of AAVs from recombinant fluids. To gather molecular-level insight in the molecular landscape of the AAV surface, we performed an *in silico* “druggability” study of the CRs using PockDrug^78,79^ and identified 5 candidate sites whose morphology and physicochemical properties are suitable for docking peptide ligands (**Figures S1B and S1C**, and **Table S1**).

The physicochemical properties of the selected sites guided the design of a peptide library for the selection of candidate ligands: *(i)* a chain length of 6 or 8 monomers was chosen based on the average size of the pockets (Van der Waals volume ∼ 950 - 1150 Å^3^; projection area ∼ 150 – 250 Å^2^), which provides an ideal balance – based on prior knowledge^50,55,58,80,81^ – between the expected biorecognition activity and manufacturing cost; *(ii)* the combinatorial positions in the library were randomized with alanine, asparagine, glutamic acid, histidine, isoleucine, lysine, phenylalanine, serine, and tryptophan, which were adopted as the amino acids capable of forming a network of diverse non-covalent interactions with the selected binding pockets; and *(iii)* a Gly-Ser-Gly (GSG) tripeptide spacer was utilized to link the combinatorial segment of the library to the resin to improve peptide display, thus promoting the outcome of library screening and subsequent Edman sequencing. The peptide libraries were synthesized following the “split-couple-and-recombine” technique^82,83^ on ChemMatrix beads – porous, hydrophilic, translucent particles that have proven an excellent substrate for the synthesis and selection of peptide ligands^57,84,85^.

The library was incubated with a model feedstock containing red-fluorescently labeled AAV2 and green-labeled HEK 293 host cell proteins (HCPs), and screened using a bead sorting device developed by our team for the rapid selection of peptide ligands^56,57^: AAV2 was utilized as model target for being the most studied and utilized of the currently known human and non-human primate AAV serotypes^23^; the formulation of the feedstock – namely AAV2 at 5·10^11^ vp/mL and HEK 293 HCPs at 0.5 mg/mL – mimics industrial cell culture lysates and was adopted to identify peptide ligands capable of isolating AAV from complex sources in *bind-and-elute* mode. The device comprises a microfluidic chamber, where each bead is imaged using a multiple wavelength fluorescence microscope, and is controlled by a software performing real-time monitoring, image processing, and selection of the beads (**Figure 1A**)^50,51,86,87^. Beads with high binding strength (*i.e.*, ratio of the bead’s *vs.* standard red fluorescence intensity > 0.9) and selectivity (*i.e.*, red *vs.* green intensity ratio > 100) are retained in the device, while all other beads are discarded. Each retained bead is exposed to a flow of elution buffer, namely 1 M MgCl_2_ in 20 mM Bis-Tris buffer at pH 6.0, whose composition and pH was adopted to ensure the selection of peptide ligands that enable efficient AAV release under gentle conditions. Accordingly, only the beads displaying effective AAV elution (*i.e.*, ratio of bead’s pre-*vs.* post-elution red fluorescence intensity > 10) were selected and analyzed via Edman degradation to sequence the peptide carried thereon. The resulting 6-mer and 8-mer sequences are listed in **Table S2**, while the homology analysis is in **Figure 1B**.

**Figure 1.**
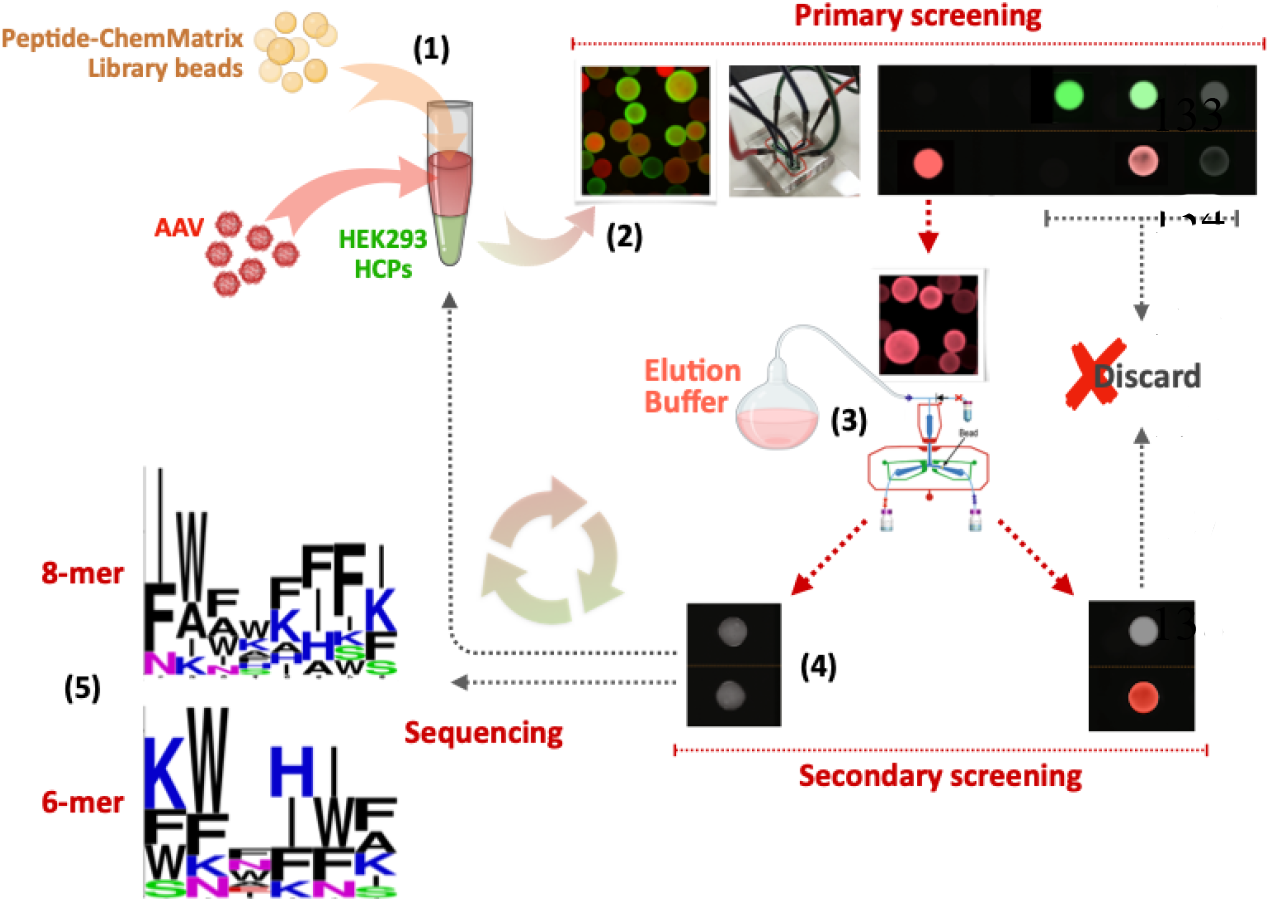
Process for identification of AAV-targeting peptide ligands. A ensemble of 6-mer or 8-mer peptide-ChemMatrix beads are (1) collectively incubated with a screening mix comprising AF594-labeled AAV2 (red) at 5·10^11^ vp/mL and AF488-labeled HE 293 HCPs (green) at ∼0.5 mg/mL; (2) the beads are fed to a microfluidic bead sorting device, which discards all non-fluorescent, the green-only, and red-and-green beads, and retains every red-only bead; (3) the latter is exposed to an elution buffer comprising 1 M MgCl_2_ in 20 mM Bis-Tris buffer at pH 6.0 for 2 mins at room temperature; (4) every bead that displays at least a 10-fold loss of red fluorescence is selected as a positive lead; (5) finally, positive beads are analyzed by Edman degradation to identify the candidate AAV-targeting peptides. (**B**) Sequence homology of the selected 6-mer and 8-mer peptides prepared using Weblogo.

### 2.2. Evaluation of AAV binding and release by peptide-functionalized chromatographic resins in non-competitive mode

Selected sequences IWWHIAKF, FWNWHHFK, FWWAAFFK, IAFKKISI, IKIFFFFS, GYISRHPG, KFNHWF, WKAHNK, KWWIWA, WWIKIS, FFNFFK, and FNHFFI were conjugated to Toyopearl AF-Amino-650M resin and evaluated for AAV binding and elution in non-competitive mode to draw an initial ranking of the candidate ligands; commercial affinity adsorbents POROS™ CaptureSelect™ AAVX and AVB Sepharose HP resin were utilized as reference standards. Since different serotypes display specific tissue tropisms^13,88^, evaluating the ability of an affinity resin to target different serotypes is the first step in demonstrating its potential in downstream bioprocessing of AAVs. In this context, we adopted AAV2 and AAV9 as model serotypes: AAV2 is the most widely studied serotype to date and is the one for which the majority of values of binding capacity of affinity adsorbents are reported in the technical literature^89–91^; AAV9 has received significant attention – clinically, for its ability to bypass the blood-brain barrier (BBB) and, in the context of biomanufacturing, for being a secreted vector whose purification is challenging (*note:* affinity resins marketed as universal AAV binders often struggle to capture AAV9, and dedicated adsorbents for AAV9 purification have been developed^13,42,44,92^). To evaluate the peptide-based resins under industrially relevant conditions, pure AAV2 and pure AAV9 at ∼5·10^11^ vp/mL in 10 mM Bis-Tris buffer at pH 7.0 were utilized as feedstocks, and loaded at a ratio of ∼10^13^ vp per mL of resin, which is expected to be the average binding capacity of the resins.

The bound AAV vectors were recovered from the peptide-Toyopearl resins under the same mild elution conditions adopted in library screening – namely, 1 M MgCl_2_ in 10 mM Bis-Tris buffer at pH 6.0 – while strong acidic elution (*i.e.*, 200 mM MgCl_2_ in 200 mM citrate buffer at pH 2.2 and PBS at pH 2.0, respectively, as recommended by the manufacturers) were implemented for the AAVX and AVB resins to ensure the most stringent performance evaluation of the selected ligands. The chromatograms obtained with pure AAV2 and AAV9 and the electrophoretic analysis of the collected fractions are respectively reported in **Figure S2** and **S3**, while the values of yield are reported in **Figure 2**.

**Figure 2.**
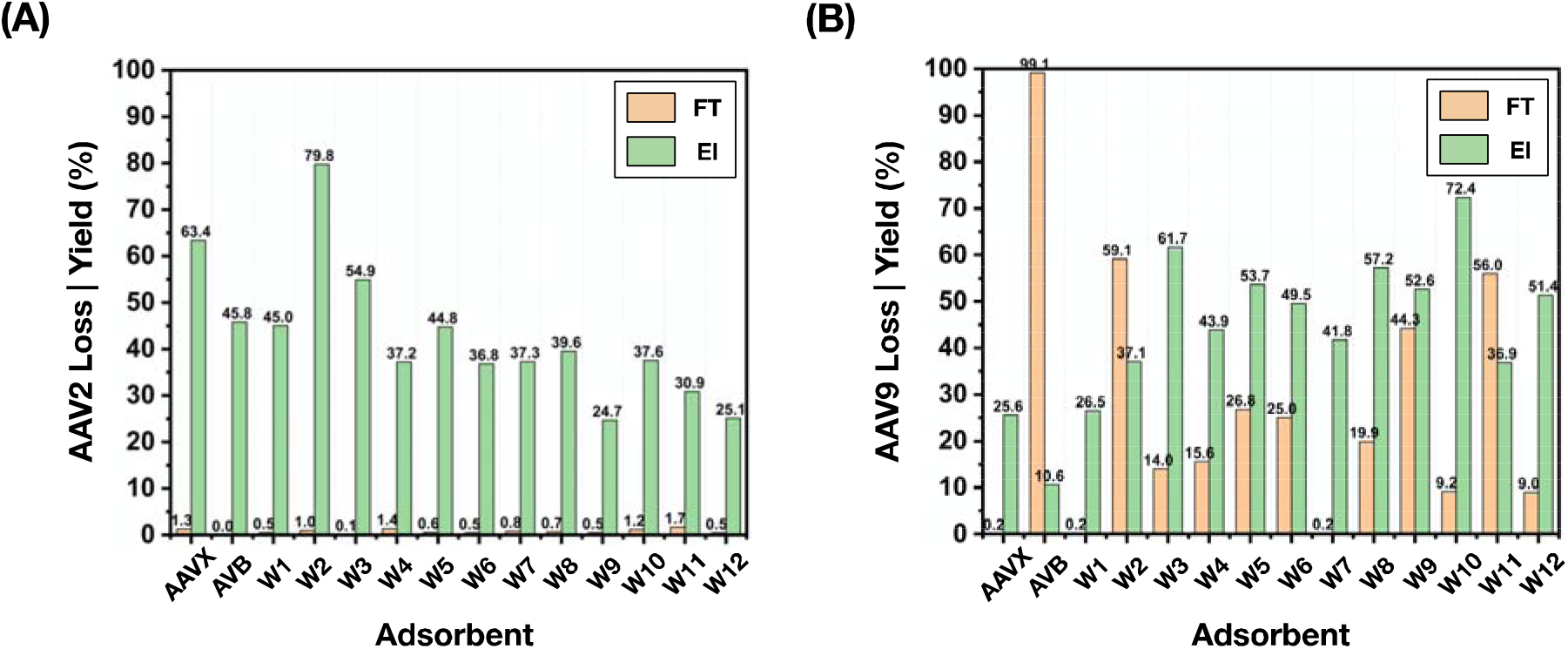
Values of loss (orange, calculated as the ratio of the AAV titer in the flow-through and wash fractions vs. load) and yield (green, calculated as the ratio of the AAV titer in the elution fraction vs. load) of (**A**) AAV2 and (**B**) AAV9 obtained via bind-and-elute studies in non-competitive mode using peptide-based resins KFNHWFG-(W1), WKAHNKG-(W2), IWWHIAKFG-(W3), FWNWHHFKG-(W4), FWWAAFFKG-(W5), IAFKKISIG-(W6), IKIFFFFSG-(W7), KWWIWAG-(W8), WWIKISG-(W9), FFNFFKG-(W10), FNHFFIG-(W11), GYISRHPG-(W12) Toyopearl resins, and control adsorbents POROS™ CaptureSelect™ AAVX and AVB Sepharose HP resins. The AAV titer in the flow-through, wash, and elution fractions was measured using serotype-specific ELISA kits.

As shown in **Figure 2**, all peptide resins bound AAV2 efficiently (< 1.7% loss). These results corroborate the library design criteria inspired by the druggability study of AAV capsid proteins and are in line with the *in-silico* evaluation of AAV: peptide binding reported below. Specifically, the values of AAV2 yield provided by KFNHWFG (45%), WKAHNKG (79.8%), IWWHIAKFG (54.9%), and FWWAAFFKG (44.8%) were comparable to those granted by AAVX POROS™ (63.4%) and AVB Sepharose (45.8%) resins. Similar values of yield were obtained with AAV9: noteworthily, IWWHIAKFG (61.7%), FWNWHHFKG (43.9%), FWWAAFFKG (53.7%), KWWIWAG (57.2%), and FFNFFKG (72.4%) significantly outperformed AAVX POROS™ (25.6%) and AVB Sepharose (10.6%) resins. Notably, while the values of yield of both serotypes were comparable across the various resins, the values of product loss in the flow-through and wash were substantially higher for AAV9 than AAV2. On the other hand, the values of yield and binding strength calculated *in silico* (K_D,*in-silico*_, see **Table 1**) indicate that product loss is not due to lack of affinity by the peptides for AAV9. The electrophoretic analysis of eluted AAV2 (**Figure S2**) and AAV9 (**Figure S3**) clearly show the presence of all three capsid proteins VP1 (∼87 kDa), VP2 (∼73 kDa), and VP3 (∼62 kDa) in the correct ∼1:1:10 ratio^1,93,94^, based on the densitometric analysis of the gels, corroborating our interpretative hypothesis that only fully formed capsids are captured by the peptide ligands.

**Table 1.** Values of dissociation constant (K_D,in silico_) of the complexes formed by peptides KFNHWFG, FFNFFKG, FNHFFIG, IWWHIAKFG, FWNWHHFKG, and FWWAAFFKG with the capsids of AAV2, AAV6, AAV8, and AAV9 obtained via molecular docking and dynamics simulations. The values of K_D,in silico_ were derived from the average of the ΔG_b_ of the various VP:peptide complexes weighted by the frequency of the interfaces, namely 3.4% for VP1-VP2, 6.9% for VP1-VP3 and VP2-VP3, and 82.8% for VP3-VP3.

| Sequence | AAV2:peptide |  | AAV6:peptide |  | AAV8:peptide |  | AAV9:peptide |  |
| --- | --- | --- | --- | --- | --- | --- | --- | --- |
| | $K_{D,in\ silico}$ (M) | | $K_{D,in\ silico}$ (M) | | $K_{D,in\ silico}$ (M) | | $K_{D,in\ silico}$ (M) | |
|  | pH 6.0 | pH 7.4 | pH 6.0 | pH 7.4 | pH 6.0 | pH 7.4 | pH 6.0 | pH 7.4 |
| KFNHWFG | $1.2 \cdot 10^{-4}$ | $1.3 \cdot 10^{-6}$ | $7.5 \cdot 10^{-4}$ | $2.0 \cdot 10^{-6}$ | $1.9 \cdot 10^{-4}$ | $2.6 \cdot 10^{-6}$ | $6.1 \cdot 10^{-5}$ | $4.6 \cdot 10^{-7}$ |
| FFNFFKG | $5.4 \cdot 10^{-4}$ | $1.1 \cdot 10^{-6}$ | $7.2 \cdot 10^{-4}$ | $3.7 \cdot 10^{-6}$ | $7.2 \cdot 10^{-4}$ | $1.0 \cdot 10^{-5}$ | $1.0 \cdot 10^{-3}$ | $1.3 \cdot 10^{-6}$ |
| FNHFFIG | $1.6 \cdot 10^{-4}$ | $9.3 \cdot 10^{-7}$ | $4.2 \cdot 10^{-4}$ | $5.2 \cdot 10^{-6}$ | $6.6 \cdot 10^{-6}$ | $1.3 \cdot 10^{-7}$ | $2.4 \cdot 10^{-3}$ | $3.7 \cdot 10^{-5}$ |
| IWWHIAKFG | $5.1 \cdot 10^{-3}$ | $2.4 \cdot 10^{-6}$ | $1.1 \cdot 10^{-3}$ | $2.7 \cdot 10^{-6}$ | $7.2 \cdot 10^{-4}$ | $2.9 \cdot 10^{-6}$ | $4.8 \cdot 10^{-3}$ | $1.1 \cdot 10^{-5}$ |
| FWNWHHFKG | $5.4 \cdot 10^{-4}$ | $1.9 \cdot 10^{-6}$ | $7.2 \cdot 10^{-4}$ | $2.4 \cdot 10^{-6}$ | $2.7 \cdot 10^{-4}$ | $5.1 \cdot 10^{-5}$ | $3.7 \cdot 10^{-3}$ | $1.3 \cdot 10^{-5}$ |
| FWWAAFFKG | $1.3 \cdot 10^{-3}$ | $1.7 \cdot 10^{-6}$ | $6.4 \cdot 10^{-4}$ | $2.3 \cdot 10^{-6}$ | $1.7 \cdot 10^{-4}$ | $8.1 \cdot 10^{-6}$ | $5.0 \cdot 10^{-3}$ | $2.5 \cdot 10^{-5}$ |

Overall, the performance of the peptide ligands is rather remarkable – with respect to their protein counterparts – when one considers the difference in elution conditions (pH 6 *vs.* pH 2). At low pH, in fact, AAVs can undergo conformational alterations that cause capsid aggregation or loss of integrity^1,95^; externalization of the VP1 phospholipase A2 (PLA_2_) domain, which triggers capsid uncoating and release of the genetic payload^1,96,97^; and biochemical alterations (*e.g.*, hydrolysis, deamidation, or oxidation of the VPs) that impair capsid docking to the cellular receptor (AAVR), and thus host entry and trafficking^98–100^. The current approach to minimizing these instances relies on immediate pH neutralization of the elution stream from POROS™ AAVX and AVB Sepharose resins. Conversely, the unique approach enabled by the peptide presented in this study aims to preserve the transduction activity of the AAV products by achieving efficient elution under near-physiological conditions

### 2.3. *In silico* investigation of AAV:peptide binding

The results presented above suggest that the identified peptide sequences are capable of binding not only the AAV2 capsids employed in the library screening, but also AAV9^101^. Notably, these two serotypes belong to different antigenic clades, respectively B and F, and feature rather different sequences (VP1/VP3 amino acid identity ∼ 81%) and structures (structural identity ∼ 94%)^61,101^.

To evaluate the ability of the selected peptides to bind multiple AAV serotypes, we modeled the binding of three 6-mer (KFNHWFG, FFNFFKG, and FNHFFIG) and three 8-mer (IWWHIAKFG, FWNWHHFKG, and FWWAAFFKG) peptides to the crystal structures of AAV2, AAV6, AAV8, and AAV9 capsids via molecular docking and molecular dynamics (MD). The secondary structure of the peptides, obtained via MD simulations in explicit water, were docked against a spherical cap representing the AAV capsids in HADDOCK v. 2.4^71,72^. The homology spherical cap structures were obtained by collating published structures, namely PDB IDs 6U0V, 6IH9, 5IPI, and 6IHB for AAV2; 3SHM, 5EGC, 4V86, 3OAH for AAV6; 6V10, 2QA0, 3RAA, 6U2V, 6PWA for AAV8; and 3UX1, 7MT0, 7WJW, and 7WJX for AAV9. An initial round of “blind” docking was performed to evaluate – in an unbiased fashion – the ability of the selected sequences to target the homologous binding sites identified in the initial “druggability” study (**Figure S1B**). Additionally, to mimic the orientational constraint imposed upon the peptides by their conjugation onto the surface of the chromatographic resin, we imposed the -GSG tripeptide appended on the C-terminal end of the peptides not to bind AAV^57,102,103,104^.

The resulting AAV:peptide complexes were refined via MD simulations (150 ns) in explicit solvent to obtain values of Gibbs free energy of binding (ΔG_b_), which were used to identify putative binding sites (|ΔG_b_| > 6.5 kcal/mol). Notably, all binding poses identified on AAV2 and AAV9 coincided with the binding sites identified in the “druggability” study. Accordingly, a second round of peptide docking was performed on these sites upon conditioning the homology structures of the capsids to both pH 7.4 and 6.0, and the docked structures were refined via extended MD simulations (500 ns) to obtain accurate values of binding energy. Representative complexes formed by the selected peptides with AAV2, AAV6, AAV8, and AAV9 capsids at pH 7.4 are shown in **Figure 3**, while the values of ΔG_b_ and the corresponding values of dissociation constant (K_D,*in*_ *_silico_*) at pH 7.4 and pH 6 are listed in **Table 1**; finally, detailed results of peptide docking for AAV2 and AAV9 are reported in **Figures S4 – S9**.

**Figure 3.**
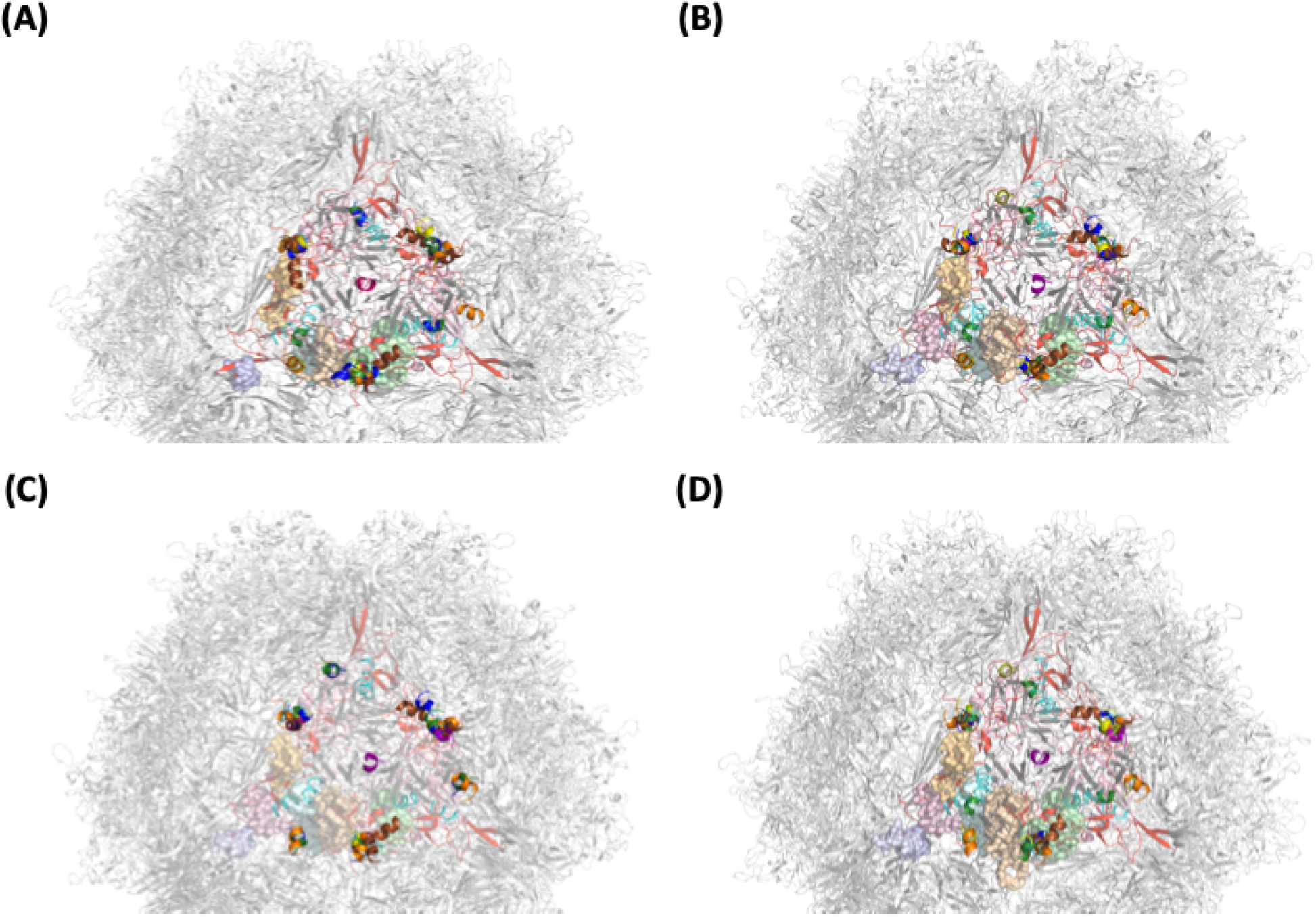
Representative complexes formed by peptides KFNHWFG (green), FFNFFKG (yellow), FNHFFIG (pink), IWWHIAKFG (brown), FWNWHHFKG (cyan), and FWWAAFFKG (orange) with the capsids of (**A**) AAV2 (PDB IDs 5IPI, 6IH9, 6IHB, and 6U0V); (**B**) AAV6 (3SHM, 3OAH, 4V86, and 5EGC); (**C**) AAV8 (2QA0, 3RAA, 6PWA, 6U2V, and 6V10); and (**D**) AAV9 (3UX1, 7MT0, 7WJW, and 7WJX) obtained via molecular docking and dynamics simulations. The segments of the VP that are not solvent accessible or whose homology among AAV serotypes is lower than 95% are in grey, the homologous segments of VP that are solvent accessible and displayed on the concave side of the capsid are in pink, and the homologous segments of VP that are solvent accessible and displayed on the convex side of the capsid are in red; the binding sites are labeled in Figure S1.

Confirming the criteria of library design and the results of dynamic binding in **Figure 2**, the *in silico* results demonstrate the ability of the selected peptides to target AAVs in a serotype-agnostic manner. Specifically, the peptides consistently conserved regions located at the interface among different VPs, which is critical in order for AAV-binding ligands to target not only multiple serotypes but also capsids of the same serotype, given the stochastic arrangement of the VPs within a capsid. Among the docked peptides, KFNHWFG and IWWHIAKFG in particular targeted homologous binding sites located at the VP1-VP2, VP1-VP3, VP2-VP3, and VP3-VP3 interfaces, while FWNWHHFKG and FNHFFIG only targeted the VP1-VP3, VP2-VP3, and VP3-VP3 interfaces. Analogous behavior is found among anti-AAV antibodies, especially those utilized in analytical and diagnostic kits^105^. As portrayed in **Figure 3**, and in more detail in **Figures S4-S9**, the pose of each peptide on homologous target sites located at different interfaces varies slightly due to subtle variations in the mutual orientation of the interlocking VPs. Because this translates in minor differences in the peptide:capsid binding energy, the values of K_D,*in*_ *_silico_* reported in **Table 2** were derived from the average of the ΔG_b_ of the various poses weighted by the frequency of the interfaces. Furthermore, peptides FNHFFIG, KFNHWFG, and IWWHIAKFG were also found to target conserved druggable domains displayed on VP3, and hence on VP1 and VP2, although they did not overlap with the binding sites of the AAV receptor (AAVR, in cyan in **Figures S4**-**S9**).

**Table 2.** Values of dynamic AAV2 binding capacity (DBC_10%_) of peptide-functionalized resins loaded with a feedstock containing AAV2 in the HEK293 lysate at the titer of 3.2·10^11^ vp/mL in PBS pH 7.4 at residence time (RT) of 3 mins.

| Resin | $DBC_{10\%}$ (vp AAV per mL of resin) |
| --- | --- |
| KFNHWF-Toyopearl | $1.33 \cdot 10^{13}$ |
| IWWHIAKF-Toyopearl | $9.53 \cdot 10^{12}$ |
| FWNWHHFK-Toyopearl | $8.75 \cdot 10^{12}$ |
| IKIFFFFS-Toyopearl | $1.02 \cdot 10^{13}$ |
| FFNFFK-Toyopearl | $1.40 \cdot 10^{13}$ |

Notably, the binding strength of the various site:peptide complexes was found to be rather weak compared to the values of the AAV:antibody counterparts (K_D,*in*_ *_silico_* ∼ 10^-9^ M). Notably, the 6-mer sequences KFNHWFG, FFNFFKG, and FNHFFIG consistently display a higher affinity, with values of K_D,*in*_ *_silico_* across the four serotypes fluctuating between 10^-6^ and 10^-7^ M, whereas 8-mers IWWHIAKFG, FWNWHHFKG, and FWWAAFFKG ranked as weaker binders, with K_D,*in*_ *_silico_*∼ 10^-5^ – 10^-6^ M. The trajectories of the molecular dynamic simulations indeed showed that the 6-mer peptides outperformed 8-mer peptides in accessing the druggable pockets located in the valleys located between the capsid’s threefold and twofold axes. Furthermore, the analysis of pairwise interactions between the residues displayed by the peptide ligands and key amino acids in the target sites of VP3 demonstrate the formation a dense network of side chain-side chain and side chain-backbone hydrogen bonds as well as π-π interactions; these account respectively for 56-68% and 11-16% of the VP:peptide binding energy; notably, fewer-than-expected coulombic and hydrophobic interactions were recorded, which provided lesser contributions, respectively 8-14% and 7-12%, to the binding energy.

Moderate binding strength is welcome in the context of affinity chromatography of labile therapeutics. For AAV purification in particular, weak VP:peptide interactions are conductive to easier elution and reduce the risk of capsid adsorption resulting in the denaturation of the protrusions that determine tissue tropism and gene transduction to the target cells. At the same time, the peptide density on the surface of the resin is sufficient to achieve multi-site interactions that grant high binding capacity and efficient product capture despite the low titer of capsids in the feedstock. Based on the values of peptide density on the resin (∼ 0.12 – 0.15 mmol per gram) and the resin’s specific surface (∼30 m^2^/g), and projection area of each asymmetric unit on the icosahedral capsid (∼81 nm^2^)^1^, approximately 30 peptides are displayed on the area of the resin that is impacted by a single capsid. Comparing this number with the arrangement of the peptide binding poses on the triangular unit shown in **Figure 3** shows the likelihood of forming 3 – 5 VP:peptide interactions per bound capsid. This ultimately suggests that AAV capture by the peptide-functionalized resin is governed by a multi-site binding mechanism, where the μM-level affinity of single peptides are synergized into nM-level avidity, on par with AAV:antibody binding. We further speculate that molecular-level non-idealities in the display of peptides or the posture of the capsid on the resin surface curb the avidity-driven capture, preventing irreversible adsorption of most of the loaded capsids; at the same time, where ideal binding arrangements occur, the mild elution conditions may fail to release the adsorbed capsids.

Following the description of the adsorption step, the *in silico* results also help elucidate AAV elution under mild conditions by offering a mechanism of capsid dissociation from the resin-bound peptides. Specifically, the molecular dynamic simulations show that the residues located at the periphery and in the immediate surrounding of the binding sites participate in a cyclical opening/closing conformational change. Decreasing the pH from 7.4 to 6.0 lowers the electrostatic charge of the binding sites by neutralizing the histidine residues (pK_a_ ∼ 6.0), the majority of which are neighbored by other cationic (*i.e.*, lysine and arginine) or anionic (*i.e.*, aspartate and glutamate); the analysis of primary sequences reported on PDB for AAV2 (6U0V), AAV6 (3SHM), AAV8 (6V10), and AAV9 (3UX1), indicates that 64% of histidine residues are either neighbored by a charged residue either immediately or with one interposing amino acid. As the local network of electrostatic bonds decrease upon acidification, the conformational flexibility of the binding sites increases, and so does the pulsatile behavior of the binding sites. As observed in prior work,^106^ oscillation and distorsion of the pairwise interactions result in lower VP:peptide binding strength: specifically, the values of |ΔG_b_| averaged over the last 100 ns of MD simulations decrease between pH 7.4 and 6.0 of ∼3.5 kcal/mol for AAV2, ∼3.3 kcal/mol for AAV6, ∼2.2 kcal/mol for AAV8, and ∼3.2 kcal/mol for AAV9. This ultimately translates in 50-200-fold variations in binding strength (K_D,*in*_ *_silico_*), which is consistent with the high values of recovery obtained experimentally. Furthermore, the addition of Mg^2+^ – a kosmotropic cation – in the elution buffer promotes a salting-in effect, leading to the hydration of the binding sites and their dissociation from the ligands. This combination of conformational pulsing and salting-in of the binding sites represents a powerful elution trigger and supports the high values of AAV yield granted by the peptide ligands.

Collectively, the *in silico* results corroborate the experimental observations that the selected sequences bind AAVs in a serotype-agnostic manner and afford efficient elution of bound capsids under mild conditions.

### 2.4. Purification of AAV from HEK 293 cell culture lysate using peptide-functionalized chromatographic resins

Having confirmed the broad targeting activity, high binding capacity, and efficient elution under mild conditions of the peptide ligands, we moved to evaluate their ability to purify AAV2 from a clarified HEK 293 cell culture lysate. The feedstock was formulated to mimic the harvests typical of the gene therapy industry (AAV2 titer ∼1.6·10^11^ vp/mL; HCP titer ∼0.5 mg/mL). A relatively short RT ∼ 3 mins was adopted for the binding step to capitalize on the high AAV binding capacity of the peptide-based adsorbents while attempting to minimize the adsorption of HEK 293 HCPs. The chromatograms of AAV2 purification are collated in **Figure S10**, while the size exclusion and steric exclusion chromatography analyses of the collected fractions are reported in **Figure S11** and **S12**, respectively; finally, the values of AAV2 yield and logarithmic removal values of HEK 293 host cell proteins (HCP LRV) are summarized in **Figure 4**.

**Figure 4.**
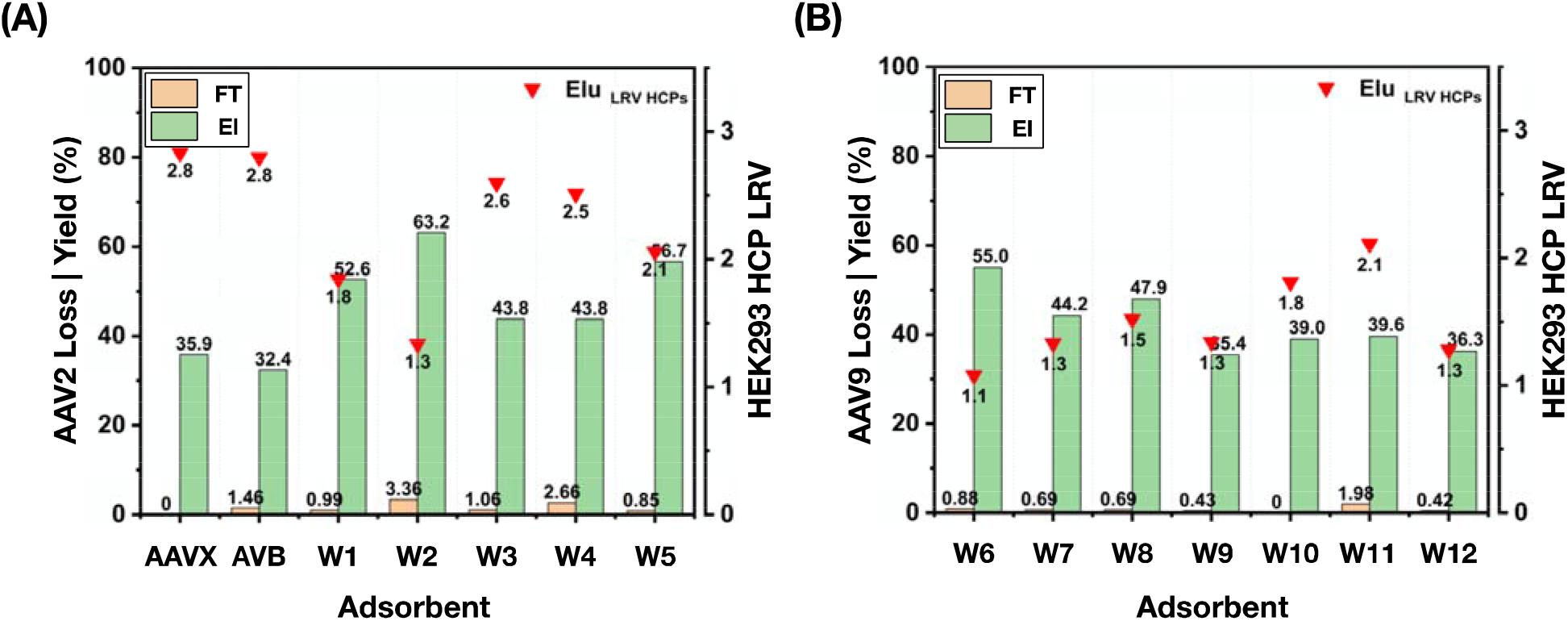
Values of loss (orange, calculated as the ratio of the AAV titer in the flow-through and wash fractions vs. load) and yield (green, calculated as the ratio of the AAV titer in the elution fraction vs. load) of AAV2 and logarithmic reduction of HCPs (HCP LRV, red triangles) obtained via chromatographic purification of AAV2 from a clarified HEK 293 cell lysate (AAV2 titer: ∼1.6·10^11^ vp/mL; HCP titer: ∼0.5 mg/mL) using (**A**) control adsorbents POROS™ CaptureSelect™ AAVX and AVB Sepharose HP resins as well as peptide-based resins KFNHWFG- (W1), WKAHNKG- (W2), IWWHIAKFG- (W3), FWNWHHFKG- (W4), FWWAAFFKG- (W5), (**B**) IAFKKISIG- (W6), IKIFFFFSG- (W7), KWWIWAG- (W8), WWIKISG- (W9), FFNFFKG- (W10), FNHFFIG- (W11), GYISRHPG- (W12) Toyopearl resins. The AAV titer in the flow-through, wash, and elution fractions was measured using serotype-specific ELISA kits.

The results in **Figure 4** mirror the values of AAV2 capture and release presented in **Figure 2A**: *(i)* product loss in the flow-through and wash fractions oscillated between 1-4 %, indicating that the adsorbents maintain their high binding capacity when loaded with complex feedstocks; *(ii)* the peptide-based adsorbents afforded yields ranging between 40-65%, thus consistently out-performing reference POROS™ CaptureSelect™ AAVX Affinity and AVB Sepharose HP resins; and *(iii)* 6 out of 12 peptide-based adsorbents afforded values of HCP LRV above 1.8, with IWWHIAKFG-Toyopearl and FWNWHHFKG-Toyopearl resins achieving LRVs of 2.6 and 2.5, respectively – corresponding to a 400-fold and 330-fold reduction of HEK 293 HCPs – thus providing a purification performance comparable to that of commercial reference resins.

The feedstock and elution fractions were further analyzed via size exclusion chromatography (SEC, **Figure S11**) and steric exclusion chromatography (SXC, **Figure S12**) to evaluate the presence of capsid aggregates and fragments, and visualize the removal of process-related impurities (*note:* while HEK 293 ELISA assays provide the titer of HCPs, analytical chromatography also reveals the presence of denatured or hydrolyzed HCPs, other non-proteinaceous metabolites, host cell DNA and RNA, and media components); and transduction assay on human epithelial cells (HT1080) to evaluate the recovery of the genetic payload and the infectivity of the purified viruses, respectively.

The SEC and SXC analyses exemplify the complex biomolecular landscape of the feedstock (**Figure S11A** and **S12A**). Mirroring the ELISA results, the chromatographic profiles of the fractions eluted from POROS™ CaptureSelect™ AAVX Affinity (**Figure S11B** and **S12B**) and AVB Sepharose HP (**Figure S11C** and **S12C**) resins highlight the high purity of the eluted AAV2; notably, despite the harsh elution pH, no aggregates were observed in those fractions, potentially owing to the salting-in effect provided by the salt composition and concentration of the respective elution buffers. The viruses eluted from the peptide-based adsorbents are accompanied by some impurities, although the chromatographic profiles of the eluted fractions from IWWHIAKFG- (∼205-fold reduction of impurities based on the chromatographic area), FWNWHHFKG- (∼160-fold), FWWAAFFKG- (185-fold), FFNFFKG- (135-fold), and FNHFFIG-Toyopearl (145-fold) resins – in line with the results of the ELISA kits – demonstrate the excellent purification activity of the selected peptide ligands.

Together with yield and purity, the transduction activity of the eluted viral vectors – namely, their ability to effectively deliver their gene payload to the target cells – is a critical parameter that defines the quality of the purification process. Unlike antibody-based therapeutics, whose biomolecular stability allows leveraging significant variations in buffer conductivity and pH to control their adsorption to and elution from the affinity resins, AAVs are significantly more susceptible to loss of activity driven by bioprocess conditions. Commercial affinity resins for AAV purification, however, mandate product elution under extremely acidic pH, varying between 2 – 3 depending upon serotype and desired elution yield (> 50%) and titer. Conversely, the peptide-based adsorbents presented in this work afford comparable elution performance under significantly milder conditions (1M MgCl_2_ in the 20 mM Bis-Tris buffer at pH 6.0). Accordingly, we resolved to quantify the transduction activity of the AAV2 purified using the peptide-based adsorbents *vs.* the reference AVB Sepharose HP, and CaptureSelect™ AAVX Affinity resins on human epithelial (HT1080) cells. The transgene encapsidated in the model AAV2 utilized in this study encodes for green fluorescence protein (GFP), thus allowing facile quantification of transduction activity via fluorescence flow cytometry. The results collated in **Figure 5** corroborate the claim that elution under mild conditions affords a product with superior transduction activity: the AAV2 purified using peptide-based adsorbents transduced in fact a remarkably higher amount of HT1080 cells, up to 20- to 60-fold more than the AAV2 eluted from commercial affinity resins.

**Figure 5.**
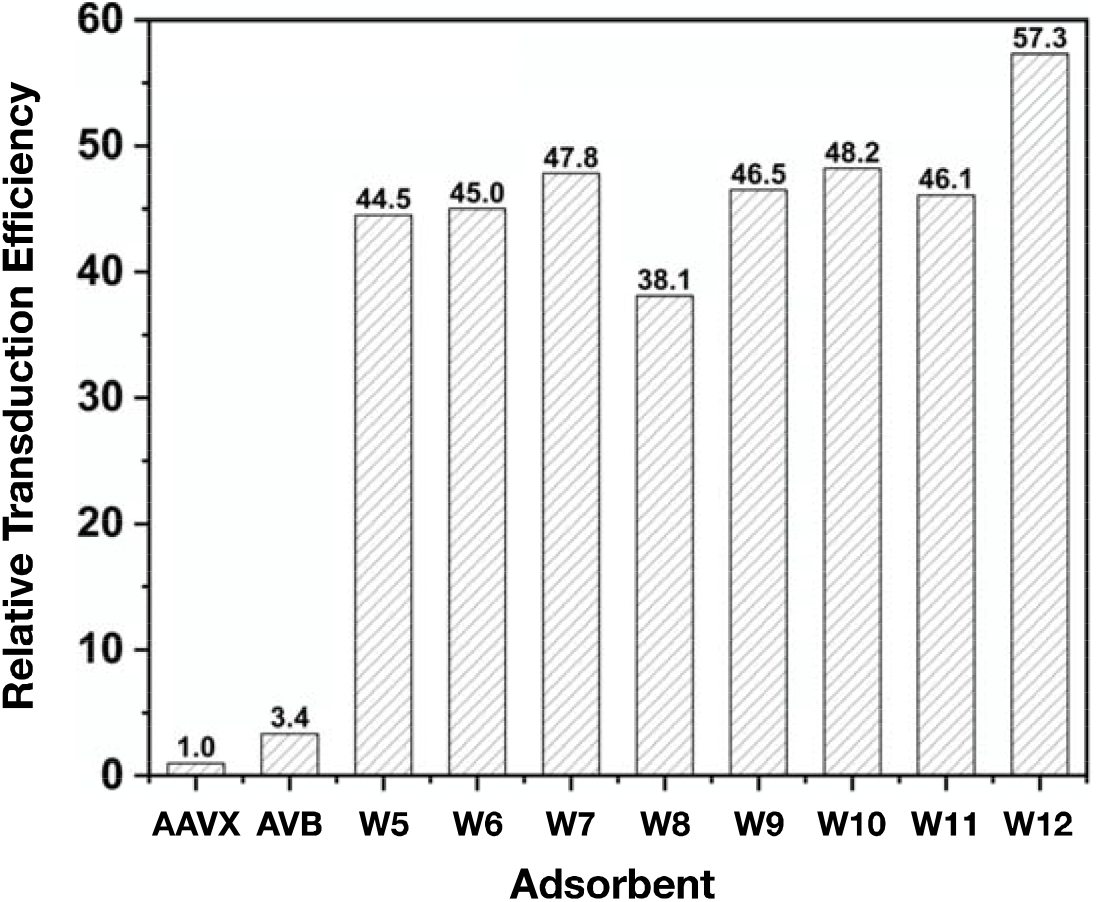
Values of relative transduction efficiency of AAV2 purified from a clarified HEK 293 cell lysate using peptide-based adsorbents FWWAAFFKG-, IAFKKISIG-, IKIFFFFSG-, KWWIWAG-, WWIKISG-, FFNFFKG-, FNHFFIG-, and GYISRHPG-Toyopearl resins-Toyopearl resins. The transduction efficiency (TU/vp) of eluted AAV2 was measured on human epithelial (HT1080) by performing a green fluorescence assay using a CytoFLEX Flow Cytometer. The values of relative transduction efficiency were calculated as the ratio of transduction efficiency of the AAV2 eluted from a peptide-based adsorbent vs. the transduction efficiency of the AAV2 eluted from POROS™ CaptureSelect™ AAVX Affinity resin.

Having compared the purification performance of the various peptides, we resolved to measure the dynamic binding capacity (DBC_10%_) of selected adsorbents KFNHWFG-, IWWHIAKFG-, FWNWHHFKG-, IKIFFFFSG-, and FFNFFKG-Toyopearl resins (**Figure S13**). All measurements were conducted at the residence time (RT) of 3 min, which is recommended for both POROS™ AAVX and AVB Sepharose resins^42,43^. The values of DBC_10%_ reported in **Table 2** met the value of 10^13^ viral particles per mL of resin – regarded by the community as the standard for affinity resins intended for AAV purification – thus supporting the adoption of peptides ligands for the product capture step in virus biomanufacturing.

## 3. Discussion

Viral vectors – and, chiefly among them, adeno-associated viruses (AAVs) – are poised to become the focus of next-generation treatment of rare cardiovascular, muscular, neurological, and ophthalmological diseases. Following the approval of LUXTURNA^®^ (AAV9) for the treatment of retinal dystrophy (2019) and ZOLGENSMA^®^ (AAV2) for treating pediatric patients with spinal muscular atrophy, late in 2022 The U.S. FDA approved CSL’s HEMGENIX^®^, an AAV5-based gene therapy for hemophilia B. With the expected uptick in regulatory approvals and the exponential growth of the clinical trials adsorbents on viral vector-based gene therapies poses a defined need for affordable tools for manufacturing AAV-based therapies, which currently carry price tags over $2.5M. In this context, a key role will be played by purification technology and particularly by affinity adsorbents for universal AAV purification. The adsorbents currently on the market have drawn substantially – both in the morphology of the beads and the design of the ligands – from the Protein A adsorbents employed in antibody purification, and are therefore poorly suited for AAV purification. Responding to these challenges, this study presented the development of the first set of serotype-agnostic AAV-targeting peptide ligands. The use of synthetic peptide ligands is particularly apt to the purification of viral vectors, since it *(i)* limits the cost of the adsorbents to below $8K per liter (*note:* this value includes the costs of the base resin, the purified peptide, and the ligand conjugation to produce 100 liters of adsorbent) against the $24-60K per liter of commercial resins; *(ii)* enables promiscuous targeting of conserved epitopes on the surface of the AAV capsids while ensuring sufficient selectivity to afford a significant reduction of process-related impurities; and *(iii)* features a moderate binding strength, thus enabling efficient elution of the bound capsids under near-physiological conditions, which are much milder than those requires for commercial affinity resins and afford an AAV product with superior transduction activity. The combination of universal serotype targeting, high binding capacity, excellent purification activity, and safeguarding the capsid’s transduction activity at a fraction of the cost of commercial adsorbents make these peptide-based adsorbents valuable tools for large scale applications supporting clinical efforts. Following on this successful demonstration, the workflow developed in this study will soon be extended to other viral vectors of medicinal relevance, such as lentivirus and adenovirus.

## 4. Materials and methods

### 4.1. Materials

Aminomethyl ChemMatrix (particle size: 100 - 200 mesh; primary amine density: 0.5 - 0.7 mmol per g) resins were sourced from PCAS Biomatrix, Inc. (Saint-Jean-sur-Richelieu, Quebec, Canada). The Toyopearl AF-Amino-650M resin (pore size: 100 nm; particle size: 65 μm; ligand density: 250 µmol per mL resin) was obtained from Tosoh Corporation (Tokyo, Japan). Fluorenylmethoxycarbonyl- (Fmoc-) protected amino acids Fmoc-Ser(tBu)-OH, Fmoc-Ile-OH, Fmoc-Phe-OH, Fmoc-Trp(Boc)-OH, Fmoc-His(Trt)-OH, Fmoc-Ala-OH, Fmoc-Asn(Trt)-OH, Fmoc-Lys(Boc)-OH, Fmoc-Glu(OtBu)-OH, and Fmoc-Gly-OH, Hexafluorophosphate Azabenzotriazole Tetramethyl Uronium (HATU), piperidine, diisopropylethylamine (DIPEA), and trifluoroacetic acid (TFA) were obtained from ChemImpex International (Wood Dale, IL, USA). Triisopropylsilane (TIPS), Kaiser test kits, 1,2-ethanedithiol (EDT), polybrene, and phosphate buffered saline (PBS) at pH 7.4 were obtained from MilliporeSigma (St. Louis, MO, USA). NHS-Alexa Fluor 594 (NHS-AF594) and NHS-Alexa Fluor 488 (NHS-AF488), N-methyl-2-pyrrolidone (NMP), N,N’-dimethylformamide (DMF), dichloromethane (DCM), methanol, POROS™ CaptureSelect™ AAVX Affinity Resin, Pluronic™ F-68 Non-ionic Surfactant, SilverQuest™ Silver Staining Kit, hydrochloric acid (HCl), sodium hydroxide (NaOH), potassium chloride (KCl), sodium chloride (NaCl), magnesium chloride (MgCl_2_), and Bis-Tris HCl were obtained from Fisher Chemical (Hampton, NH, USA). Dulbecco’s Modified Eagle Medium (DMEM) and fetal bovine serum (FBS) were sourced from ThermoFisher (Waltham, MA). Human fibrosarcoma (HT1080) cells were obtained from ATCC (Manassas, VA). The AVB Sepharose HP was sourced from Cytiva (Marl-borough, MA). Clarified HEK 293 cell lysates containing AAV2, and null HEK 293 cell culture lysate were donated by BTEC (Raleigh, NC, USA). All chromatographic experiments were performed using a ÄKTA Avant system from Cytiva (Marlborough, MA, USA). Alltech chromatography columns (diameter: 3.6 mm; length: 50 mm; volume: 0.5 mL), and 10 µm polyethylene frits were obtained from VWR International (Radnor, PA, USA). A BioResolve SEC mAb Column (particle diameter: 2.5 µm; pore diameter: 200Å; column diameter: 7.8 mm; column length: 300 mm) size exclusion chromatography column was obtained from Waters Inc. (Milford, MA, USA). A CIMac PrimaS^™^ 0.1 mL analytical monolith column (diameter: 5.2 mm; length: 4.95 mm; volume: 0.1 mL, channel radius: 1050 nm) for steric exclusion chromatography analysis was obtained from BIA separations (Ajdovscina, Slovenia). The 10-20% Tris-Glycine HCl SDS-PAGE gels were purchased from Bio Rad Life Sciences (Hercules, CA, USA). The AAV ELISA kits were purchased from Progen (Wayne, PA, USA) while the HEK 293 ELISA kits were purchased from Cygnus (Southport, NC, USA).

### 4.2. Synthesis of peptide libraries on aminomethyl-ChemMatrix resin and selected peptides on Toyopearl resin

Peptide synthesis was performed on a Syro I automated peptide synthesizer (Biotage, Uppsala, Sweden) using nine protected amino acids, namely Fmoc-Ser(tBu)-OH, Fmoc-Ile-OH, Fmoc-Phe-OH, Fmoc-Trp(Boc)-OH, Fmoc-His(Trt)-OH, Fmoc-Ala-OH, Fmoc-Asn(Trt)-OH, Fmoc-Lys(Boc)-OH, and Fmoc-Glu(OtBu)-OH.^50,51^ Each amino acid coupling step was performed at 45°C for 20 min, using 3 equivalents (eq.) of protected amino acid at the concentration of 0.5 M, 3 eq. of HATU and 0.5 M, 6 eq. of DIPEA (0.5M) in 5 mL of dry DMF. The yield of peptide conjugation was monitored after each amino acid via the Kaiser test. The removal of Fmoc protecting groups was performed at room temperature using 20% piperidine in DMF. The 6-mer (X_1_-X_2_-X_3_-X_4_-X_5_-X_6_) and 8-mer (X_1_-X_2_-X_3_-X_4_-X_5_-X_6_-X_7_-X_8_) peptide libraries were synthesized on 2 g of Aminomethyl-ChemMatrix resin preloaded with the tripeptide spacer GSG (G: glycine; S: serine) following the “split-couple-recombine” method^50,52–54^. Selected peptides IWWHIAKF, FWNWHHFK, FWWAAFFK, IAFKKISI, IKIFFFFS, GYISRHPG, KFNHWF, WKAHNK, KWWIWA, WWIKIS, FFNFFK, and FNHFFI were synthesized on Toyopearl AF-Amino-650M resin at the density of ∼ 0.12 - 0.15 mmol of peptide per gram of resin (*note:* minor sequence-dependent variability in peptide density is expected). Upon completing chain elongation, the peptides were deprotected via acidolysis using a cleavage cocktail containing TFA, thioanisole, anisole, and EDT (94/3/2/1) for 2 hrs. After deprotection, the ChemMatrix library resins were rinsed with DCM and DMF and stored in DMF at 4°C, whereas the peptide-Toyopearl resins were washed sequentially with DCM, DMF, methanol, and stored in 20% v/v aqueous methanol.

### 4.3. Fluorescent Labeling of AAV and HEK 293 HCPs

The labeling method of viral vector and proteins was performed as reported in the previous studies^50,55^. The AAV2, AAV9 were labeled using NHS-Alexafluor 594 (NHS-AF594, red), while the host cell proteins (HCPs) contained in the HEK 293 cell culture lysate were collectively labeled with NHS-Alexafluor 488 (NHS-AF488, green). Both dyes were initially dissolved in anhydrous DMSO to a concentration of 10 mg/mL. A volume of 1 μL of NHS-AF594 was added to 100 μL of AAV solution at 10^13^ vp/mL in PBS pH 7.4, while 200 μL of NHS-AF488 was added to 4 mL of null HEK 293 cell culture lysate at the HCP titer of 300 μg/mL. The labeling reactions were allowed to proceed for 1 hr at room temperature, under dark and gentle agitation. The unreacted dyes were removed using 0.5 mL Zeba™ Dye and Biotin Removal Spin Columns (ThermoFisher Scientific, Waltham, MA). The concentration of the labeled AAV2 and AAV9 in solution was determined by AAV ELISA assay and the concentration of the labeled proteins in solution was determined by Bradford assay. The absorbance of the solutions of AF488-labeled HEK 293 HCPs and AF594-labeled AAV was measured by UV spectrophotometry at the wavelength of 490 and 590 nm, respectively, using a Synergy H1 plate reader (Biotek, Winooski, VT).

### 4.4. Dual-Fluorescence Screening of Peptide Library against AAV in HEK 293 cell culture lysate

The screening process of AAV-targeting peptides was performed as reported in the previous studies^50,55^. A screening mix was initially prepared by spiking AF594-labeled AAV2 and AAV9 in AF488-labeled HEK 293 cell culture lysate to obtain a final AAV titer of 5·10^11^ vp/mL and a HEK 293 HCP titer of ∼0.5 mg/mL. Aliquots of 10 μL of library beads were initially equilibrated with PBS at pH 7.4 and subsequently incubated with 40 μL of screening mix for 2 hrs at room temperature in dark. The beads were thoroughly washed with PBS at pH 7.4 and 0.1% v/v Tween 20 in PBS at pH 7.4 and sorted automatically using the microfluidic screening device developed in prior work^55–57^. The beads were fed to microfluidic sorter, which is controlled by a custom-made MATLAB code programmed to discard the non-fluorescent, the green-only, and red-and-green beads, and withhold beads with strong red-only fluorescence. The latter were exposed to a flow of 1 M MgCl_2_ in 20 mM Bis-Tris HCl buffer at pH 6.0 for 2 mins at room temperature. Every bead that displayed a measurable loss of red fluorescence was selected as a positive lead, individually incubated with 100 μL of 0.1 M glycine buffer at pH 2.5 for 1 hr at room temperature in the dark to elute the bound AF594-labeled AAV, and rinsed with Milli-Q water and acetonitrile. The beads were finally analyzed via Edman degradation using a PPSQ-33A protein sequencer (Shimadzu, Kyoto, Japan) to identify the peptide sequences carried thereupon.

### 4.5. Dynamic binding capacity of peptide-Toyopearl resins

The dynamic binding capacity at 10% breakthrough (DBC_10%_, vp per mL resin) of peptide-Toyopearl resins were measured as reported in prior studies^50,58,59^. A volume of 0.5 mL of resin packed in an Alltech chromatography column was initially washed with 10 column volumes (CVs) of 20% v/v ethanol and deionized water (3 CVs) and equilibrated with 10 CVs of PBS buffer. A volume of 30 mL of solution of AAV2 at 3.2·10^11^ vp/mL in HEK 293 lysate (HCP titer ∼ 0.5 mg/mL) was loaded on the column at the flow rate of 0.17 mL/min (residence time, RT: 3 min). The resin was then washed with 20 CVs of binding buffer at 0.5 mL/min. The AAV elution from the peptide-Toyopearl resins was performed using 1 M MgCl_2_ in 20 mM Bis-Tris HCl buffer at pH 6.0 at the flow rate of 0.25 mL/min (RT: 2 min); as recommended by manufacturers, elution from POROS™ CaptureSelect™ AAVX affinity resin and AVB Sepharose HP resin was performed at the flow rate of 0.25 mL/min (RT: 2 min) using 0.2 M MgCl_2_ in 200 mM citrate buffer at pH 2.2 and PBS at pH 2.0, respectively. All resins were regenerated with 10 CVs of PBS buffer at pH 2.0 at the flow rate of 0.5 mL/min. The effluents from flowthrough step were continuously monitored by UV spectrometry at 280 nm and collected in volume of 1 mL per tube. The collected fractions were analyzed by AAV ELISA Kit to generate the breakthrough curves, and the resulting chromatograms were utilized to calculate the DBC_10%_.

### 4.6. Purification of AAV from HEK 293 cell culture lysate using peptide-Toyopearl resins

Each peptide-functionalized resin and commercial resins was individually wet packed in the 0.5 mL Alltech column, washed with 20% v/v ethanol (10 CVs) and deionized water (3 CVs), and equilibrated with 10 CVs of 20 mM NaCl in 20 mM Bis-Tris HCl buffer at pH 7.0. A volume of 12 mL of AAV2 at ∼1.6·10^11^ vp/mL in HEK 293 cell culture lysate (HCP titer ∼ 0.5 mg/mL) was loaded on the column at the flow rate of 0.17 mL/min (residence time, RT: 3 min). Following load, the resin was washed with binding buffer (20 CVs) to recover the UV_280nm_ baseline. The AAV elution from the peptide-Toyopearl resins was performed using 1 M MgCl_2_ in 20 mM BisTris HCl buffer at pH 6.0 at the flow rate of 0.25 mL/min (RT: 2 min); elution from POROS™ CaptureSelect™ AAVX affinity resin and AVB Sepharose HP resin was performed at the flow rate of 0.25 mL/min (RT: 2 min) using 0.2 M MgCl_2_ in 200 mM citrate buffer at pH 2.2 and PBS at pH 2.0, respectively. All resins were regenerated with 10 CVs of PBS buffer at pH 2.0 at the flow rate of 0.5 mL/min. The collected flow-through and elution fractions were analyzed by AAV2 ELISA Kit (see *Section 2.7*) to measure the AAV titration and determine the values of product yield, and HEK 293 ELISA Kit (see *Section 2.8*), size exclusion chromatography (see *Section 2.9*), steric exclusion chromatography (see *Section 2.10*), gel electrophoresis (see *Section 2.11*) to determine the values of product purity, and fluorescence flow cytometry (see *Section 2.12*) to quantify the transduction efficiency of the eluted AAVs.

### 4.7. Quantification of AAV yield

The AAV titer in the feedstock, flow-through, and elution fractions collected as described in *Section 2.5* and *Section 2.6* was measured using a AAV Titration ELISA kit (PROGEN, Wayne, PA) following the manufacturer’s protocol. The total capsids of AAV adsorbed per volume of resin was calculated via mass balance.

### 4.8. Quantification of HEK 293 HCPs

The titer of HEK 293 HCPs in the feedstock, flowthrough and elution fractions collected as described in *Section 2.6* were determined using a HEK 293 HCP ELISA kit (Cygnus Technologies, Southport, NC) following the manufacturer’s protocol. The values of HCP LRV in the effluents were calculated via mass balance.

### 4.9. Qualification of AAV purity by size-exclusion chromatography (SEC)

The feedstock, and the flow-through and elution fractions collected as described in *Section 2.6* were analyzed by analytical SEC using a BioResolve SEC mAb Column (Waters, Milford, MA) operated with a 40-min isocratic method using 200 mM KCl in 50 mM sodium phosphate at pH 7.0 (0.05% v/v sodium azide) as mobile phase. A volume of 10 µL of sample was injected and the effluent continuously monitored via UV (abs: 260 nm and 280 nm) fluorescence spectroscopy (ex/em: 280/350 nm).

### 4.10. Qualification of AAV purity by steric-exclusion chromatography (SXC)

The feedstock, the flow-through, and elution fractions collected as described in *Section 2.6* were analyzed by analytical SXC using a monolith CIMac PrimaS™ 0.1 mL analytical column (BIA Separations, Slovenia) operated with a 20-min linear gradient from 100:0 A:B to 0:100 A:B at the flow rate of 0.33 mL/min (mobile phase A: PBS at pH 7.0 added with 10% v/v PEG 6K; mobile phase B: 3X PBS at pH 7.0). Injection volumes were normalized via AAV ELISA titer to compare each elution condition and the effluent continuously monitored via fluorescence spectroscopy (ex/em: 280/350 nm).

### 4.11. Quantification of AAV purity by sodium dodecyl sulfate polyacrylamide gel electrophoresis (SDS-PAGE)

The feedstock, the flow-through, and elution fractions collected as described in *Section 2.6* were analyzed by reducing SDS-PAGE using 4-20% Mini-PROTEAN^TM^ TGX^™^ Precast protein gels (Bio Rad) and 1X Tris/Glycine/SDS Buffer (Bio Rad) as running buffer. A volume of 40 µL of sample, each adjusted to a total AAV titer of ∼1·10^12^ vp/mL, was loaded to the wells of SDS-PAGE gels. The sample stripes were concentrated under 80 V for about 30 min and separated under 120 V for about 1 hr. The gels were then stained using a SilverQuest™ Silver Staining Kit (ThermoFisher, Waltham, MA) and finally imagined by the Gel Doc2000 imaging system (Bio Rad).

### 4.12. Quantification of AAV transduction efficiency via fluorescence flow cytometry (FFC)

HT1080 cells were cultured in DMEM media supplemented with 10% v/v FBS at 5% CO_2_ and 37°C. Upon reaching 80-90% confluence, cells were seeded in 96-well plates at a density of 6,000 cells/well and cultured overnight. The eluted fractions containing AAVs were serially diluted in DMEM medium (without FBS and antibiotics) added with polybrene at 8 μg/mL, and 0.1 mL of diluted AAV sample was incubated with the HT1080 cells in the 96-well plates. After 24 hrs, spent medium was replaced with fresh DMEM containing FBS and the cells were cultured for 3 days. The percentage of cells expressing GFP (GFP^+^) was quantified using a CytoFlex flow cytometer (Beckman Coulter, Brea, CA) and the number of transduction units per mL (TU/mL) was calculated using Equation 1:

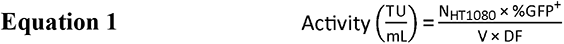

Wherein *N_HT1080_* is the number of cells incubated with each diluted AAV sample, *V* is the volume of the diluted AAV sample, and *DF* is the dilution factor.

### 4.13. In silico identification of putative peptide binding sites on AAV and analysis of AAV: peptide complexes

The crystal structures of AAV2 and AAV9 were initially prepared using Protein Prep Wizard (PPW, Schrödinger, New York, NY)^60^ by correcting missing atoms and side chains, removing salt ions, adding explicit hydrogens, and optimizing the hydrogen-bonding network. Four clusters, each comprising 15 virion proteins – namely, one VP1, one VP2, and thirteen VP3 to recapitulate the ∼1:1:10 ratio reported for AAV capsids^13,61^ – and arranged in a spherical cap that represents ¼ of the capsid surface were obtained from the Protein DataBank for AAV2 (PDB IDs: 6U0V, 6IH9, 5IPI, and 6IHB) and AAV9 (3UX1, 7MT0, 7WJW, and 7WJX). The ionization states at pH 7.4 and pH 6.0, and the corresponding structural minimization of the eight VP clusters were performed using PROPKA^62,63^. The adjusted structure was then analyzed using SiteMap to identify sites for peptide binding^64–66^, and the sites with high S-score (> 0.8) and D-score (> 0.9) were selected for peptide docking. The AAV2:antibody complex (3J1S) and the AAV9:AAVR complex (7WJX) were utilized as reference structures. Peptides IWWHIAKF, FWNWHHFK, FWWAAFFK, IAFKKISI, IKIFFFFS, GYISRHPG, KFNHWF, WKAHNK, KWWIWA, WWIKIS, FFNFFK, and FNHFFI were constructed using the molecular editor Avogadro^67^ and their structures were simulated in GROMACS using the GROMOS 43a1 force field. Briefly, each peptide was initially placed in a simulation box with periodic boundary containing 1,000 TIP3P water molecules and equilibrated with 10,000 steps of steepest gradient descent; and heated to 300 K in an NVT ensemble for 250 ps using 1 fs time steps, and equilibrated to 1 atm via a 500-ps NPT simulation with 2 fs time steps. Production runs were then performed in the NPT ensemble at 300 K and 1 atm using the Nosé-Hoover thermostat and the Parrinello-Rahman barostat^68,69^; the leap-frog algorithm with integration steps of 2 fs was used to integrate the motion equations; the covalent bonds were constrained using the LINCS algorithm, the Lennard-Jones and shortLJrange electrostatic interactions were calculated using cut-off values of 1.0 nm and 1.4 nm, respectively, while the particle-mesh Ewald method was utilized for long-range electrostatic interactions^70^; the lists of bonded and non-bonded interactions (cutoff of 1.4 nm) were updated every 2 fs and 5 fs, respectively. The energetic landscape associated to the various peptide conformations was sampled to identify the structures with absolute energy minima. The selected peptide ligands were then docked in silico against the putative binding sites on AAV2 and AAV9 using the docking software HADDOCK (High Ambiguity Driven Protein-Protein Docking, v.2.4)^71,72^. The residues on the selected binding sites of AAV2 and AAV9, and the X_1_-X_2_-X_3_-X_4_-X_5_-X_6_(-X_7_-X_8_) residues on the peptides were marked as “active”, while the surrounding residues were marked as “passive”. The docked AAV:peptide structures were grouped in clusters of up to 20 complexes based on Cα RMSD < 7.5 Å and ranked using the dMM-PBSA score^73^. Finally, the top AAV:peptide complexes were refined via 150-ns atomistic MD simulations and evaluated to estimate the free energy of binding (ΔG_B_).

## Acknowledgements

The authors wish to acknowledge the funding provided by the National Science Foundation (CBET 1743404 and CBET 1653590), the Novo Foundation (AIM-Bio Grant NNF19SA0035474) as well as the generous support of the Golden LEAF Biomanufacturing Training and Education Center (BTEC) and the North Carolina Viral Vector Initiative in Research and Learning (NC-VVIRAL) at NC State University.

## Author Contributions

W.C., S.S., E.B., R.P, P.G.-K., W.S., B.M., R.K., and C.C. conducted the experimental work. W.C., J.P., G.G, M.A.D., and S.M. conceived the work and wrote the manuscript.

