## Supplementary information for "Peptide ligands for the affinity purification of adeno-associated viruses from HEK 293 cell lysates"

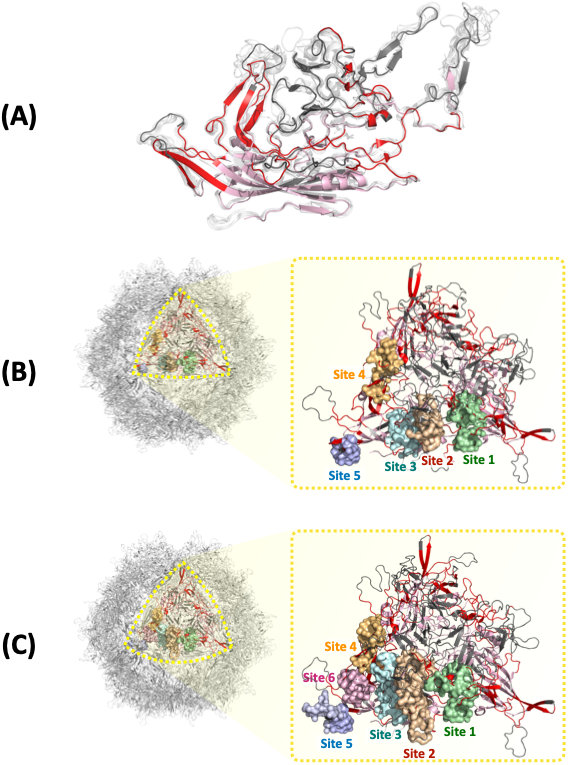

***Figure S1. Druggability study of AAV virion protein (VP). (A)*** *Structure of the VP1 from AAV1 (PDB ID: 6JCR), AAV2 (6IH9), AAV3 (3KIC), AAV4 (2G8G), AAV5 (7KP3), AAV6 (5EGC), AAV7 (7JOT), AAV8 (2QA0), and AAV9 (7WJX); the segments of the VP that are not solvent accessible or whose homology among AAV serotypes is lower than 95% are in grey cartoon, the homologous segments of VP that are solvent accessible and displayed on the concave side of the capsid are in pink cartoon, and the homologous segments of VP that are solvent accessible and displayed on the convex side of the capsid are in red cartoon;* ***(B)*** *putative binding sites 1 – 5 identified on the homologous segments of VP that are solvent accessible and displayed on the convex side of the AAV2 capsid; and* ***(C)*** *putative binding sites 1 – 6 identified on the homologous segments of VP that are solvent accessible and displayed on the convex side of the AAV9 capsid.*

***Figure S2.*** ***(A) – (C)*** *Chromatograms of AAV2 binding and elution using peptide-based adsorbents KFNHWFG- (W1), WKAHNKG- (W2), IWWHIAKFG- (W3), FWNWHHFKG- (W4), FWWAAFFKG- (W5), IAFKKISIG- (W6), IKIFFFFSG- (W7), KWWIWAG- (W8), WWIKISG- (W9), FFNFFKG- (W10), FNHFFIG- (W11), GYISRHPG- (W12) Toyopearl resins, and control adsorbents POROS™ CaptureSelect™ AAVX Affinity and AVB Sepharose HP resins. Binding was conducted in 20 mM NaCl in 20 mM Bis-Tris buffer at pH 7.0 (RT: 3 min); elution from the peptide-functionalized resins was conducted using 1M MgCl_2_ in 20 mM Bis-Tris buffer at pH 6.0 (RT: 2 min); elution from POROS™ CaptureSelect™ AAVX affinity resin and AVB Sepharose HP resin was conducted using 0.2 M MgCl_2_ in 200 mM citrate buffer at pH 2.2 and PBS at pH 2.0, respectively (RT: 2 min). (D) SDS-PAGE analysis (reducing condition, silver staining) of the elution fractions; labels: MW, molecular weight marker; AAV2 standard; VP, virion proteins; E, eluted fraction.*

***Figure S3.*** ***(A) – (C)*** *Chromatograms of AAV9 binding and elution using peptide-based adsorbents KFNHWFG- (W1), WKAHNKG- (W2), IWWHIAKFG- (W3), FWNWHHFKG- (W4), FWWAAFFKG- (W5), IAFKKISIG- (W6), IKIFFFFSG- (W7), KWWIWAG- (W8), WWIKISG- (W9), FFNFFKG- (W10), FNHFFIG- (W11), GYISRHPG- (W12) Toyopearl resins, and control adsorbents POROS™ CaptureSelect™ AAVX Affinity and AVB Sepharose HP resins. Binding was conducted in 20 mM NaCl in 20 mM Bis-Tris buffer at pH 7.0 (RT: 3 min); elution from the peptide-functionalized resins was conducted using 1M MgCl_2_ in 20 mM Bis-Tris buffer at pH 6.0 (RT: 2 min); elution from POROS™ CaptureSelect™ AAVX affinity resin and AVB Sepharose HP resin was conducted using 0.2 M MgCl_2_ in 200 mM citrate buffer at pH 2.2 and PBS at pH 2.0, respectively (RT: 2 min). (D) SDS-PAGE analysis (reducing condition, silver staining) of the elution fractions; labels: MW, molecular weight marker; AAV9 standard; VP, virion proteins; E, eluted fraction.*

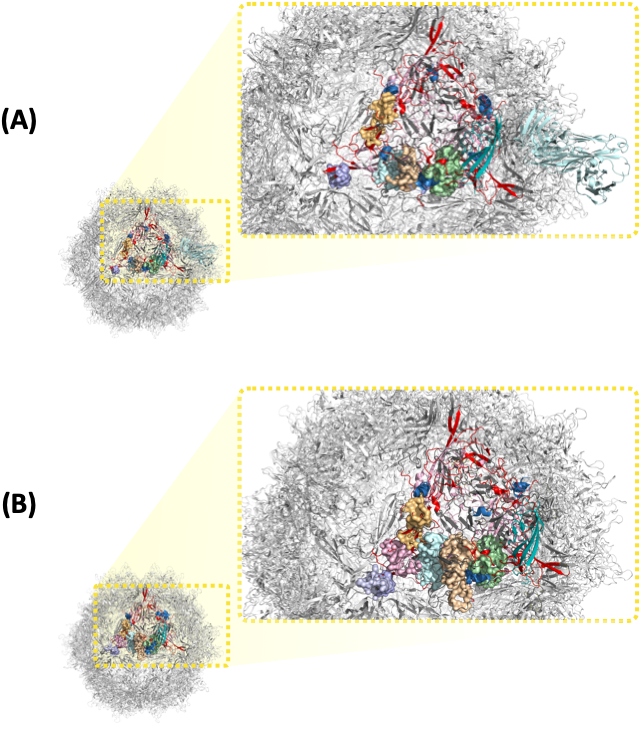

***Figure S4.*** *Complexes formed by peptide KFNHWF-GSG with the solvent accessible peptide segments displayed on the convex side of the of VP1-VP2-VP3 cluster of AAV2 (PDB IDs: 6U0V, 6IH9, 5IPI, and 6IHB) and AAV9 (3UX1, 7MT0, 7WJW, and 7WJX). The segments of the VP that are not solvent accessible or whose homology among AAV serotypes is lower than 95% are in grey cartoon, the homologous segments of VP that are solvent accessible and displayed on the concave side of the capsid are in pink cartoon, and the homologous segments of VP that are solvent accessible and displayed on the convex side of the capsid are in red cartoon; the binding sites are labeled in* ***Figure S1****; the peptide ligands are in blue cartoon.*

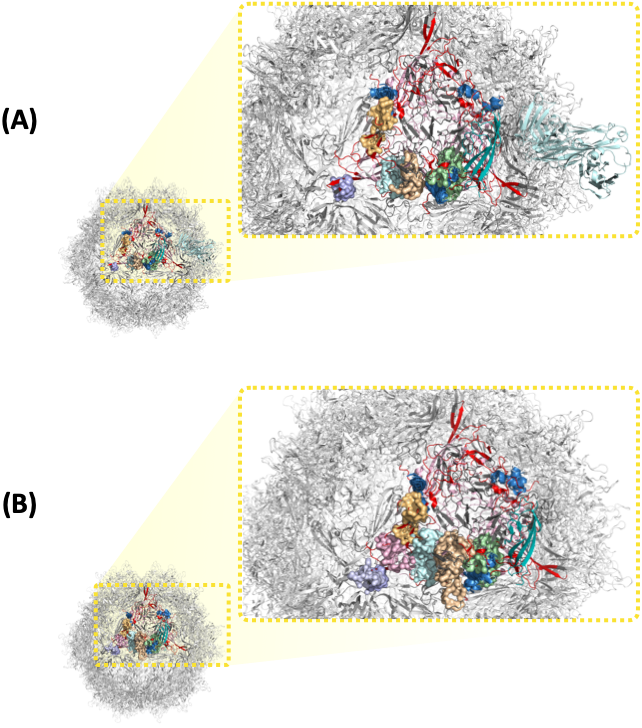

***Figure S5.*** *Complexes formed by peptide IWWHIAKF-GSG with the solvent accessible peptide segments displayed on the convex side of the of VP1-VP2-VP3 cluster of AAV2 (PDB IDs: 6U0V, 6IH9, 5IPI, and 6IHB) and AAV9 (3UX1, 7MT0, 7WJW, and 7WJX). The segments of the VP that are not solvent accessible or whose homology among AAV serotypes is lower than 95% are in grey cartoon, the homologous segments of VP that are solvent accessible and displayed on the concave side of the capsid are in pink cartoon, and the homologous segments of VP that are solvent accessible and displayed on the convex side of the capsid are in red cartoon; the binding sites are labeled in* ***Figure S1****; the peptide ligands are in blue cartoon.*

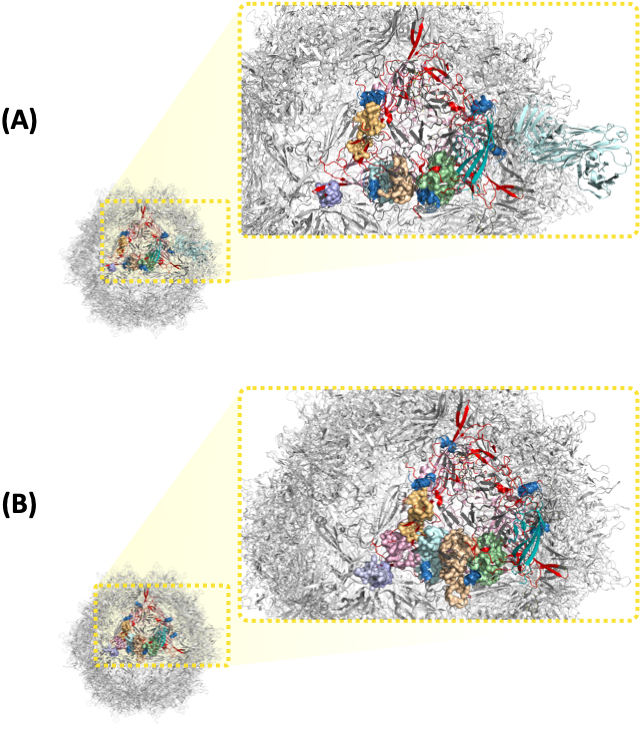

***Figure S6.*** *Complexes formed by peptide FWWAAFFK-GSG with the solvent accessible peptide segments displayed on the convex side of the of VP1-VP2-VP3 cluster of AAV2 (PDB IDs: 6U0V, 6IH9, 5IPI, and 6IHB) and AAV9 (3UX1, 7MT0, 7WJW, and 7WJX). The segments of the VP that are not solvent accessible or whose homology among AAV serotypes is lower than 95% are in grey cartoon, the homologous segments of VP that are solvent accessible and displayed on the concave side of the capsid are in pink cartoon, and the homologous segments of VP that are solvent accessible and displayed on the convex side of the capsid are in red cartoon; the binding sites are labeled in* ***Figure S1****; the peptide ligands are in blue cartoon.*

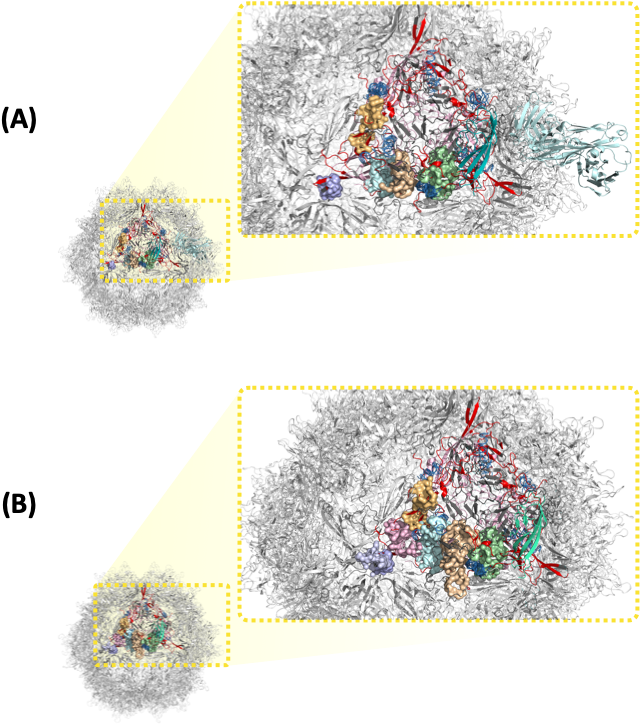

***Figure S7.*** *Complexes formed by peptide FWNWHHFK-GSG with the solvent accessible peptide segments displayed on the convex side of the of VP1-VP2-VP3 cluster of AAV2 (PDB IDs: 6U0V, 6IH9, 5IPI, and 6IHB) and AAV9 (3UX1, 7MT0, 7WJW, and 7WJX). The segments of the VP that are not solvent accessible or whose homology among AAV serotypes is lower than 95% are in grey cartoon, the homologous segments of VP that are solvent accessible and displayed on the concave side of the capsid are in pink cartoon, and the homologous segments of VP that are solvent accessible and displayed on the convex side of the capsid are in red cartoon; the binding sites are labeled in* ***Figure S1****; the peptide ligands are in blue cartoon.*

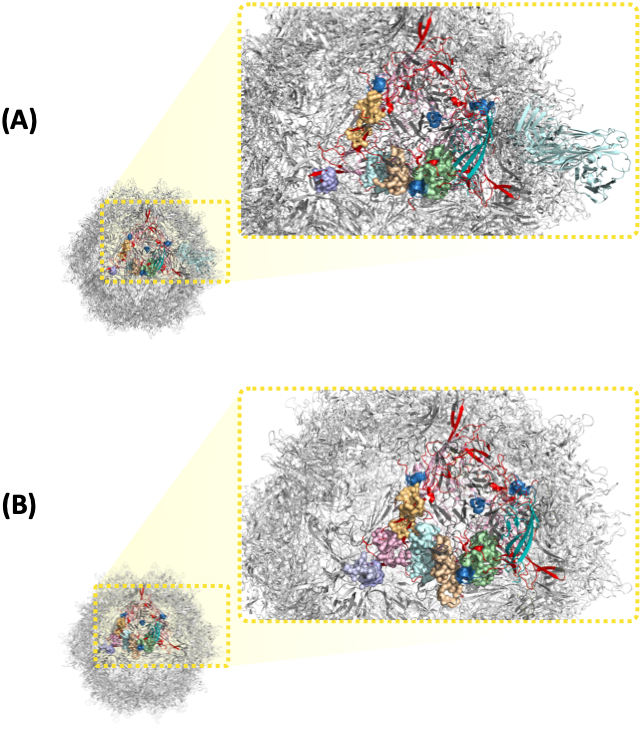

***Figure S8.*** *Complexes formed by peptide FNHFFI-GSG with the solvent accessible peptide segments displayed on the convex side of the of VP1-VP2-VP3 cluster of AAV2 (PDB IDs: 6U0V, 6IH9, 5IPI, and 6IHB) and AAV9 (3UX1, 7MT0, 7WJW, and 7WJX). The segments of the VP that are not solvent accessible or whose homology among AAV serotypes is lower than 95% are in grey cartoon, the homologous segments of VP that are solvent accessible and displayed on the concave side of the capsid are in pink cartoon, and the homologous segments of VP that are solvent accessible and displayed on the convex side of the capsid are in red cartoon; the binding sites are labeled in* ***Figure S1****; the peptide ligands are in blue cartoon.*

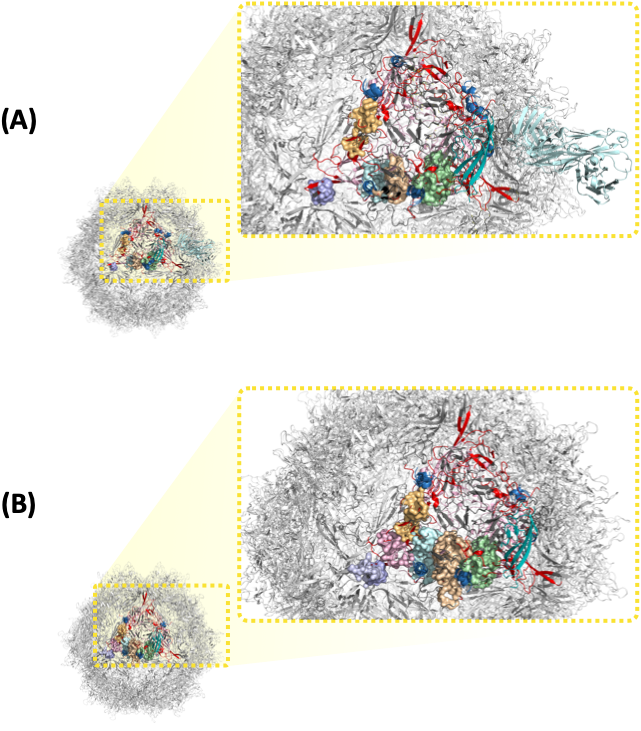

***Figure S9.*** *Complexes formed by peptide FFNFFK-GSG with the solvent accessible peptide segments displayed on the convex side of the of VP1-VP2-VP3 cluster of AAV2 (PDB IDs: 6U0V, 6IH9, 5IPI, and 6IHB) and AAV9 (3UX1, 7MT0, 7WJW, and 7WJX). The segments of the VP that are not solvent accessible or whose homology among AAV serotypes is lower than 95% are in grey cartoon, the homologous segments of VP that are solvent accessible and displayed on the concave side of the capsid are in pink cartoon, and the homologous segments of VP that are solvent accessible and displayed on the convex side of the capsid are in red cartoon; the binding sites are labeled in* ***Figure S1****; the peptide ligands are in blue cartoon.*

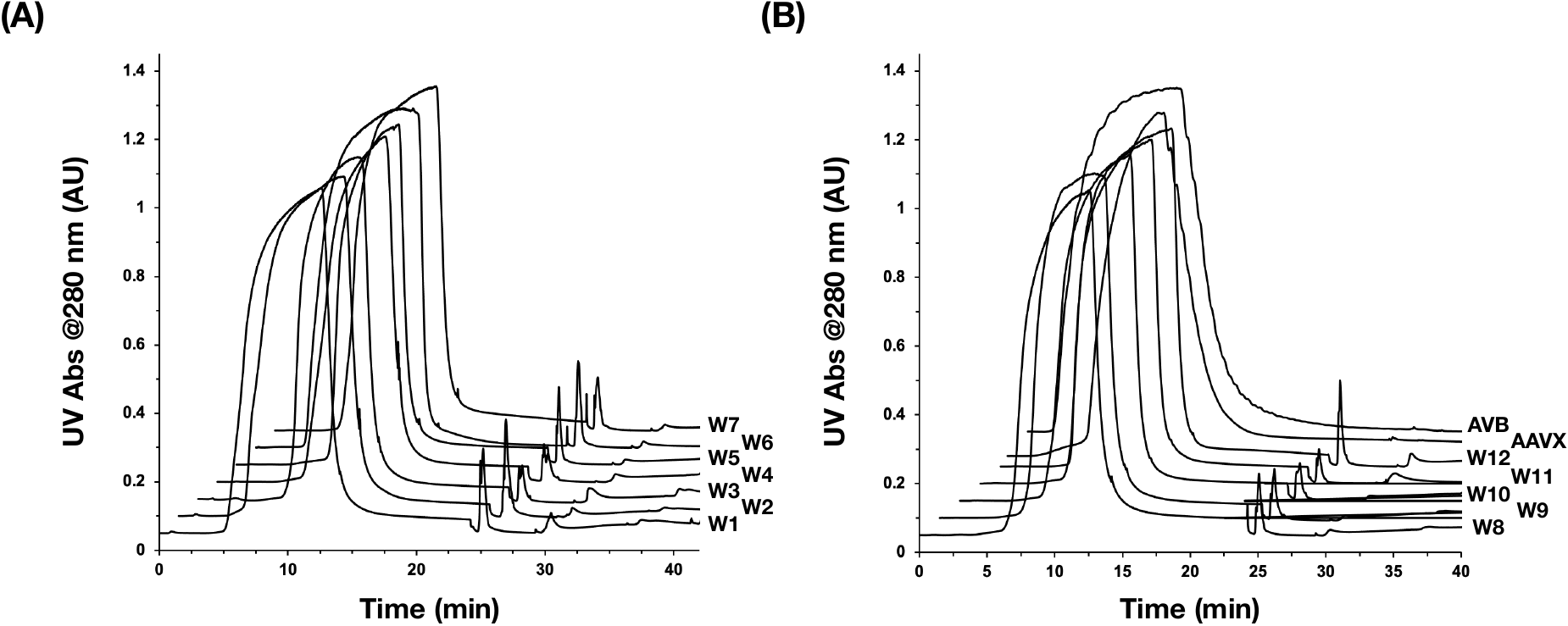

***Figure S10.*** ***(A) – (C)*** *Chromatograms of AAV2 purification from a clarified HEK 293 cell lysate (AAV2 titer: ~1.6·10^11^ vp/mL; HCP titer: ~0.5 mg/mL) using peptide-based adsorbents KFNHWFG- (W1), WKAHNKG- (W2), IWWHIAKFG- (W3), FWNWHHFKG- (W4), FWWAAFFKG- (W5), IAFKKISIG- (W6), IKIFFFFSG- (W7), KWWIWAG- (W8), WWIKISG- (W9), FFNFFKG- (W10), FNHFFIG- (W11), GYISRHPG- (W12) Toyopearl resins, and control adsorbents POROS™ CaptureSelect™ AAVX and AVB Sepharose HP resins. Binding was conducted in 20 mM NaCl in 20 mM Bis-Tris buffer at pH 7.0 (RT: 3 min); elution from the peptide-functionalized resins was conducted using 1M MgCl_2_ in 20 mM Bis-Tris buffer at pH 6.0 (RT: 2 min); elution from POROS™ CaptureSelect™ AAVX affinity resin and AVB Sepharose HP resin was conducted using 0.2 M MgCl_2_ in 200 mM citrate buffer at pH 2.2 and PBS at pH 2.0, respectively (RT: 2 min).*

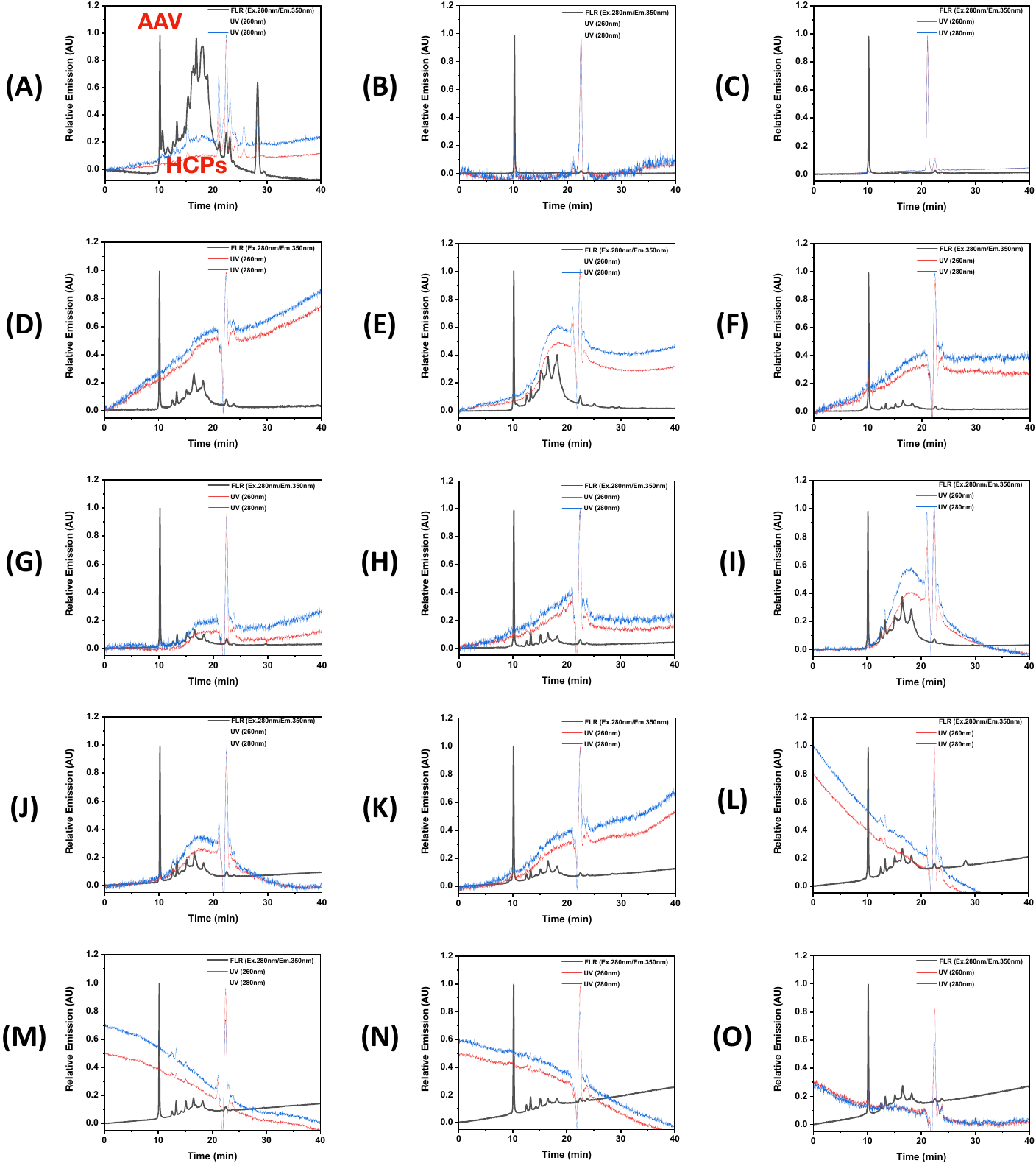

***Figure S11.*** *Size Exclusion Chromatography (SEC) analysis of* ***(A)*** *a clarified HEK 293 cell lysate (AAV2 titer: ~1.6·10^11^ vp/mL; HCP titer: ~0.5 mg/mL) and the elution fractions obtained from the purification of AAV2 from the clarified lysate using control* ***(B)*** *POROS™ CaptureSelect™ AAVX Affinity resin and* ***(C)*** *AVB Sepharose HP resin as well as peptide-based adsorbents* ***(D)*** *KFNHWFG-,* ***(E)*** *WKAHNKG-,* ***(F)*** *IWWHIAKFG-,* ***(G)*** *FWNWHHFKG-,* ***(H)*** *FWWAAFFKG,* ***(I)*** *IAFKKISIG-,* ***(J)*** *IKIFFFFSG-,* ***(K)*** *KWWIWAG-,* ***(L)*** *WWIKISG-,* ***(M)*** *FFNFFKG-,* ***(N)*** *FNHFFIG-, and* ***(O)*** *GYISRHPG-Toyopearl resins.*

***Figure S12.*** *Steric Exclusion Chromatography (SXC) analysis of* ***(A)*** *a clarified HEK 293 cell lysate (AAV2 titer: ~1.6·10^11^ vp/mL; HCP titer: ~0.5 mg/mL) and the elution fractions obtained from the purification of AAV2 from the clarified lysate using control* ***(B)*** *POROS™ CaptureSelect™ AAVX Affinity resin and* ***(C)*** *AVB Sepharose HP resin as well as peptide-based adsorbents* ***(D)*** *KFNHWFG-,* ***(E)*** *WKAHNKG-,* ***(F)*** *IWWHIAKFG-,* ***(G)*** *FWNWHHFKG-,* ***(H)*** *FWWAAFFKG,* ***(I)*** *IAFKKISIG-,* ***(J)*** *IKIFFFFSG-,* ***(K)*** *KWWIWAG-,* ***(L)*** *WWIKISG-,* ***(M)*** *FFNFFKG-,* ***(N)*** *FNHFFIG-, and* ***(O)*** *GYISRHPG-Toyopearl resins.*

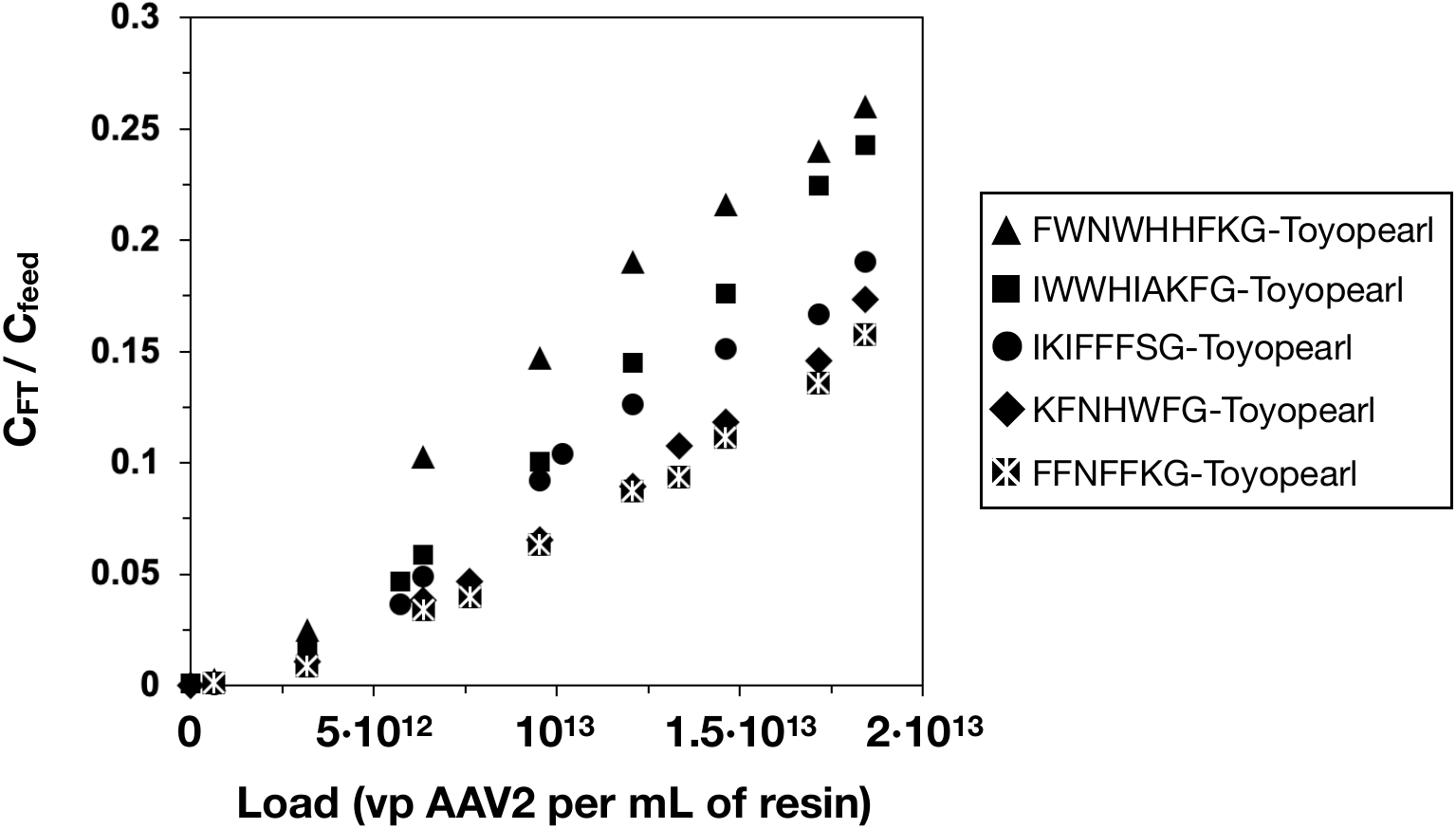

***Figure S13****. Breakthrough curves of AAV2 obtained by loading a clarified HEK 293 cell lysate (AAV2 titer: ~3.2·10^11^ vp/mL; HCP titer: ~0.5 mg/mL) on adsorbents KFNHWFG-, IWWHIAKFG-, FWNWHHFKG-, IKIFFFFSG-, and FFNFFKG-Toyopearl resins at residence time (RT) of 3 mins.*

***Table S1. Structural and biophysical properties of target sites on AAV Capsids.*** *Properties of the putative binding sites identified via SiteMap on the homologous, solvent accessible peptide segments displayed on the convex side of the of VP1-VP2-VP3 cluster of AAV2 (PDB ID: 6IH9) and AAV9 (7WJX); the binding sites are labeled in* ***Figure S1****.*

| **AAV2** | | | | | | |
| --- | --- | --- | --- | --- | --- | --- |
| **Site** | **SAS (Å^2^)** | **Volume (Å^3^)** | **Ip** | **Gravy** | **Polarity** | **Score** |
| 1 | 265.7 | 1953.9 | 8.23 | -0.605 | 0.583 | 0.892 |
| 2 | 290.3 | 2832.5 | 7.85 | -0.543 | 0.645 | 0.890 |
| 3 | 277.0 | 2079.8 | 7.48 | -0.470 | 0.470 | 0.833 |
| 4 | 295.7 | 2121.7 | 8.47 | -0.403 | 0.663 | 0.907 |
| 5 | 154.9 | 869.8 | 4.63 | -0.335 | 0.585 | 0.743 |
| **AAV9** | | | | | | |
| **Site** | **SAS (Å^2^)** | **Volume (Å^3^)** | **Ip** | **Gravy** | **Polarity** | **Score** |
| 1 | 214.9 | 1917.4 | 8.72 | -0.668 | 0.564 | 0.862 |
| 2 | 348.4 | 2703.6 | 7.64 | -0.612 | 0.636 | 0.937 |
| 3 | 322.6 | 1907.2 | 7.27 | -0.628 | 0.434 | 0.812 |
| 4 | 270.1 | 1689.1 | 8.27 | -0.610 | 0.573 | 0.927 |
| 5 | 176.9 | 969.8 | 4.55 | -0.393 | 0.537 | 0.723 |
| 6 | 182.2 | 951.2 | 6.72 | -0.487 | 0.343 | 0.862 |

***Table S2. Sequences and biophysical properties of selected 6-mer and 8-mer AAV-binding peptide ligands.*** *The values of isoelectric point (pI), polarity (Grantham scale), and Grand Average Hydropathy index (GRAVY) were calculated based on the amino acid sequence and assuming an amidated C-terminus to represent the conjugation of the peptide to the chromatographic resin.*

| **Sequence** | **pI** | **Polarity** | **GRAVY** |  | **Sequence** | **pI** | **Polarity** | **GRAVY** |
| --- | --- | --- | --- | --- | --- | --- | --- | --- |
| FFNFFK | 10.13 | 5.65 | -1.13 |  | FWNWHHFK | 11.9 | 7.95 | -1.20 |
| SWFIIF | 9.01 | 5.47 | -1.95 |  | IWAWFHFF | 10.9 | 7.86 | -2.14 |
| KWFIIF | 11.13 | 5.88 | -1.50 |  | FWWAAFFK | 10.2 | 7.55 | -1.54 |
| FNEFFI | 6.99 | 5.22 | -1.02 |  | IWWHIAKF | 10.9 | 7.44 | -1.36 |
| WKAHNK | 11.66 | 5.03 | 0.3 |  | IKIFFFFS | 11.8 | 7.04 | -1.29 |
| WWIKIS | 11.03 | 5.64 | -1.18 |  | NWAWFIWK | 11.8 | 7.81 | -1.48 |
| KFNHWF | 10.93 | 5.78 | -0.95 |  | FIFSKFFI | 11.5 | 7.03 | -1.29 |
| KWWIWA | 11.23 | 6.18 | -1.58 |  | IAFKKISI | 12.1 | 6.02 | -0.26 |
| KWWHWA | 10.83 | 6.24 | -1.37 |  | IAFKKIII | 11.7 | 6.33 | -0.53 |
